# Toward understanding drivers of specialized metabolism in Actinomycetota: insights from 1432 transcriptomics datasets for 132 strains

**DOI:** 10.64898/2026.06.17.732583

**Authors:** Anna Rudenko, Omkar Mohite, Boin Yun, Byung Tae Lee, Byeongsub Lee, Joon Young Kwon, Hahk-Soo Kang, Alberto Santos, Tilmann Weber, Hyun Uk Kim, Pep Charusanti

**Affiliations:** The Novo Nordisk Foundation Center for Biosustainability, Technical University of Denmark, Kgs. Lyngby, 2800, Denmark; Department of Biotechnology and Biomedicine, Technical University of Denmark, Kgs. Lyngby, 2800, Denmark; Department of Biomedical Science and Engineering, Konkuk University, Gwangjin-gu, Seoul, 05029, South Korea; Department of Chemical and Biomolecular Engineering, Korea Advanced Institute of Science and Technology, Daejeon, 34141, South Korea; Graduate School of Engineering Biology, Korea Advanced Institute of Science and Technology, Daejeon, 34141, South Korea; Department of Health Technology, Technical University of Denmark, Kgs. Lyngby, 2800, Denmark

**Author notes:** Contact Information: Anna Rudenko, Omkar Mohite, Boin Yun, Byung Tae Lee, Byeongsub Lee, Joon Young Kwon, Hahk-Soo Kang, Alberto Santos, Tilmann Weber, Hyun Uk Kim, Pep Charusanti.

**Keywords:** Multi-Omics, Transcriptomics, Actinobacteria, Specialized Metabolism, Gene Regulation

## Abstract

Actinomycetota are major sources of specialized metabolites with applications in drug discovery and agriculture, yet much of their biosynthetic potential remains silent under standard laboratory conditions. Limited understanding of the mechanisms controlling biosynthetic gene cluster (BGC) activation constrains metabolite discovery and production. Here, we generated 1432 RNA-seq datasets from 132 Actinomycetota strains grown in eight media to identify patterns associated with BGC expression. On average, strains expressed 44% of their encoded BGCs across the tested conditions, with more expression observed among known BGCs (61%) compared to uncharacterized BGCs (35%). Co-expression analyses revealed frequent associations between BGCs and transporters, transcriptional regulators, and proteins containing DUF397 and DUF742 domains. Targeted overexpression of candidate genes selected from BGC-associated co-expression modules increased metabolite production, with DUF397- and DUF742-containing operons showing the broadest effects by boosting the levels of several different specialized metabolites. Other genes boosted levels in a metabolite-specific manner. Together, our results support a multilayered model of BGC regulation in Actinomycetota in which BGC expression is shaped by medium composition, BGC-specific regulators, and integration of BGCs into broader transcriptional network modules. By connecting BGC expression to specific media and co-expressed genes, this study also provides a resource for selecting growth conditions and engineering specialized metabolism.

## BACKGROUND

*Streptomyces* are prolific producers of secondary (specialized) metabolites, including antibiotics, antifungals, immunosuppressants, and antitumor agents [1, 2]. Their genomes typically encode 25–50 biosynthetic gene clusters (BGCs), contiguous sets of genes required for specialized metabolite biosynthesis, but approximately 80% to 90% are silent or cryptic under commonly used laboratory conditions [3]. The large proportion of silent BGCs reflects the tight and complex regulation of BGC expression, which depends on specific environmental signals such as nutrient limitation, stress, or interspecies interactions that are not recapitulated in standard cultivation conditions [4].

BGC expression is controlled by multilayered regulatory networks [4]. Pathway-specific regulators located within BGCs often directly activate biosynthetic gene expression [5, 6], but their activity is frequently modulated by global regulators. For example, the PhoR–PhoP two-component system responds to phosphate limitation and can induce antibiotic production [7, 8]; DasR links N-acetylglucosamine metabolism to the repression or activation of multiple antibiotic pathways [9]; and BldA couples morphological differentiation to antibiotic production [10]. Additional pleiotropic regulators, including AtrA, AdpA, and Crp, also influence specialized metabolism across multiple loci [11, 12]. More recently, conserved XRE–DUF397 operons and conservon-associated DUF742-domain proteins have emerged as additional global regulatory elements that can modulate antibiotic production and activate cryptic pathways [13–16].

Although these studies have provided important insight into BGC regulation, their scope is inherently limited because they often focus on individual regulators, specific pathways, or single strains. An alternate approach is to analyze large-scale datasets spanning multiple conditions or strains to identify recurring patterns that are not readily apparent from smaller, more focused studies. In specialized metabolism, this approach has revealed variation in BGC diversity and precursor availability across different strains and microbial natural product chemical diversity across large genomic and metabolomic datasets [17–20]. Similarly, DNA Affinity Purification Sequencing (DAP-seq) studies have identified regulatory connections between transcription factors (TFs) and BGCs by mapping the binding sites for multiple TFs throughout the genome [21–23].

Transcriptomics is well suited to resolve BGC regulation because co-expression patterns can reveal co-expression modules linking biosynthetic genes to candidate regulators. Applied to large RNA-seq compendia, iModulon analyses have begun to elucidate transcriptional regulatory networks in *Streptomyces coelicolor* and *Streptomyces albidoflavus* [24–26]. Co-expression analyses in fungi have revealed novel associations between biosynthetic genes and regulators located outside BGCs [26]. However, apart from iModulons, which typically require >100 RNA-seq datasets of the same strain cultivated under different conditions, most transcriptomic studies in Actinomycetota have been limited to small numbers of strains and few conditions, providing context-specific insights but limited generalizability [28–30].

To address this gap, we generated 1432 RNA-seq datasets from 132 Actinomycetota strains profiled across multiple media conditions, with most strains (n = 117) analyzed across eight conditions comprising five rich media (DNPM, ISP2, MA, SoyM50, TSB50) and three defined minimal media (SMM-gluc, SMM-gly, SMM-malt), and smaller subsets across seven (n = 8) or up to fourteen (n = 7) conditions. The strains are all part of the Technical University of Denmark Natural Bioactive Compound (NBC) collection [31]. We then developed an integrated data analysis framework combining genome mining, large-scale transcriptomics, and weighted co-expression network analysis (WGCNA) [32, 33] (Fig. 1). This approach enables identification of recurring transcriptional patterns associated with BGC activation, including co-expression modules enriched for BGC genes and candidate regulators such as hypothetical proteins and DUF families. Experimental validation of selected candidates demonstrates that perturbing these nodes can enhance the production of multiple specialized metabolites, revealing new features of the regulatory network governing specialized metabolism and suggesting new strategies to engineer Actinomycetota strains.

**Fig. 1.**
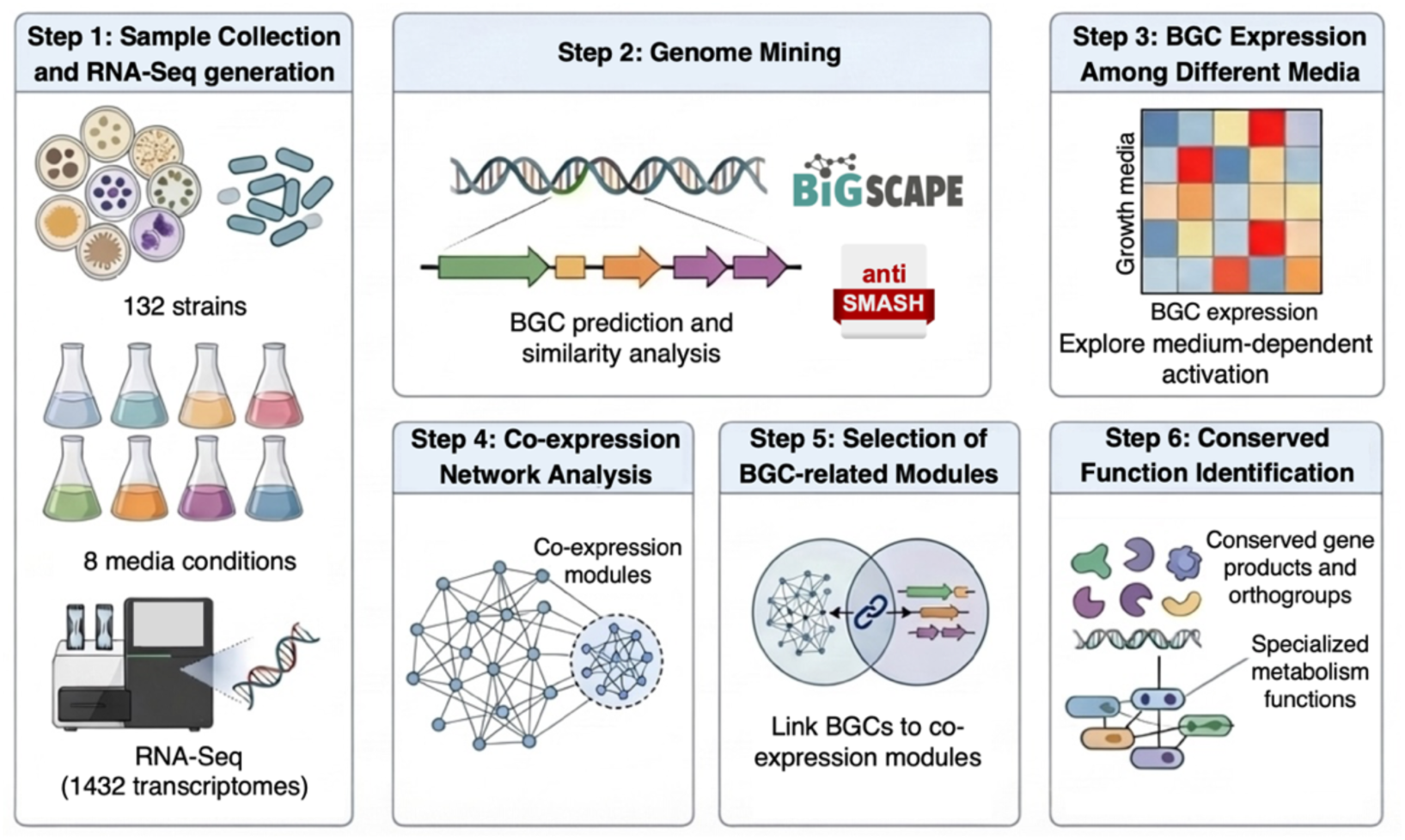
Integrated multi-omics framework to identify genes commonly co-expressed with BGCs. RNA-seq was performed across 132 strains grown under multiple media conditions. Co-expression network analysis and integration with genomic data (antiSMASH, BiG-SCAPE) enabled BGCs to be linked to transcriptional modules. Subsequently, conserved gene functions associated with specialized metabolism were identified throughout the genome based on genome-wide co-expression patterns.

## RESULTS

### BGC expression patterns across eight media

We first queried the complete RNA-Seq dataset to estimate the number of BGCs that were expressed. BGCs were determined in each genome by use of antiSMASH 7.0 [34]. To estimate transcriptional activation, we applied a per-sample dynamic threshold: for each RNA-Seq library, we computed the 75^th^ percentile of gene expression values across all genes and defined a BGC as expressed if > 50% of its core biosynthetic genes exceeded this threshold, highlighting highly expressed BGCs with a greater likelihood of producing specialized metabolites. Given that antiSMASH predictions do not accurately predict BGC boundaries, we focused our analysis only on core biosynthetic genes. For sensitivity analysis, we also report results using a more permissive 50^th^ percentile threshold. Additionally, we compared expression patterns between known and unknown BGCs. BGCs were classified as “known” if at least 80% of their genes matched a characterized cluster in the MiBiG database [35]. All remaining BGCs were classified as unknown.

Across all strains, there were 3993 antiSMASH-predicted BGCs in total, with the fraction classified as expressed showing a broad distribution (Supplementary Fig. 1). Using the stringent 75^th^-percentile rule, strains expressed on average 43.7% of their BGCs (IQR = 36.7–50.0%, range = 19.2–70.0%). Under the more permissive 50th-percentile rule, the expressed fraction increased to a mean of 62.9% (IQR = 57.1–69.9%, range = 26.9–95.0%). A total of 29 strains showed expression of more than half of their BGCs at the 75^th^ percentile threshold, increasing to 117 strains at the 50^th^ percentile threshold. Detailed information on BGC-specific expression and silent clusters for each strain is provided in Supplementary Table 1 for the 50^th^ percentile and Supplementary Table 2 for the 75^th^ percentile. For all downstream analyses, we applied the 75^th^ percentile threshold to define BGC expression. To retain expression-level resolution beyond binary expressed/not-expressed calls, we also examined the distribution of mean core biosynthetic gene expression across BGC types and media (Supplementary Fig. 2).

There were 1046 known BGCs, of which 61% (641) were expressed. In contrast, only 35% (1028) of 2947 unknown BGCs were expressed. On average, known BGCs were expressed in four media conditions compared to three for unknown BGCs. Among the eight tested conditions, the highest number of expressed clusters was seen in MA, DNPM, and SoyM50, with 870, 837 and 813 expressed BGCs, respectively (Fig. 2A). Fewer expressed BGCs were seen in ISP2 and TSB50 (755 and 616, respectively). Across all media, the majority of expressed BGC types were annotated as other (192–233 per medium) or hybrid (122–192 per medium), followed by terpene (127–169 per medium) and siderophore (31–76 per medium).

**Fig. 2.**
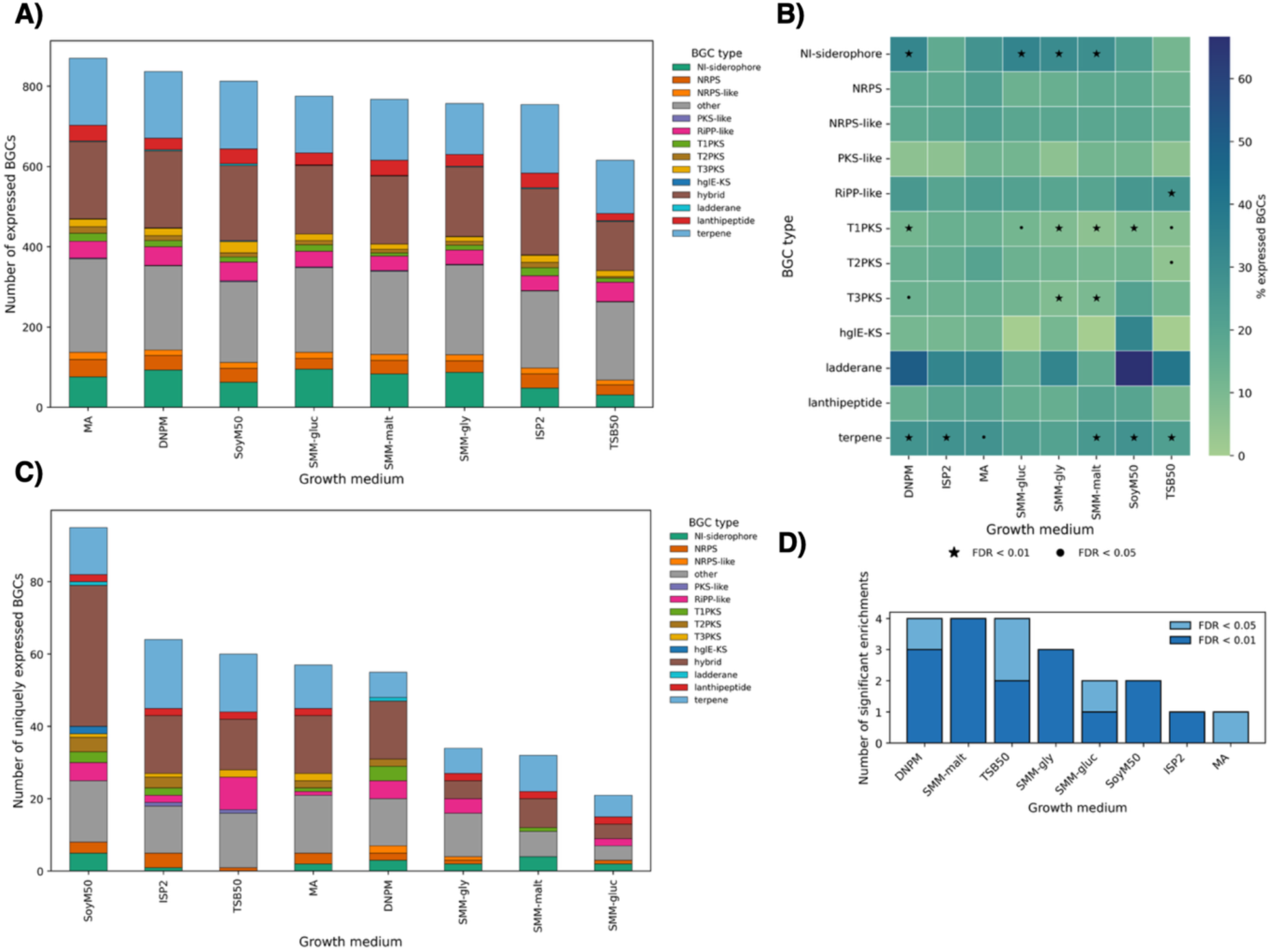
Medium-dependent expression of BGCs. **A)** Total number and composition of expressed BGCs in each media. **B)** Heatmap showing the percentage of expressed BGCs for each cluster type and medium. Asterisks indicate statistically significant enrichment of BGC activation (Fisher’s exact test, Benjamini–Hochberg correction; black circle FDR < 0.05, black star FDR < 0.01). **C)** Total number and composition of uniquely expressed BGCs in each media. **D)** Number of BGC types significantly enriched among expressed BGCs in each growth medium. Bars indicate the number of BGC classes with significant medium-specific enrichment, with colors showing enrichments detected at FDR < 0.05 and FDR < 0.01.

To investigate whether specific media were associated with the expression of otherwise silent clusters, we identified BGCs whose expression was restricted to a single medium and that remained transcriptionally inactive across all other tested media (Fig. 2C). SoyM50 supported the highest number of medium-specific BGC expression events, with 95 BGCs expressed only in this condition, dominated by the hybrid (39) and other (17) categories. ISP2 and TSB50 each activated 64 and 60 medium-specific BGCs. The three defined media resulted in only 20 to 29 medium-specific activations, mainly representing terpene and “other” BGCs.

To systematically assess whether specific media preferentially activate certain classes of BGCs, we applied Fisher’s exact tests to all media and BGC-type combinations (Fig. 2B,D). For each medium, the test compared the fraction of expressed clusters of a given type to the fraction of expressed clusters of all other types, evaluating whether that BGC class was significantly over- or under-represented among the expressed clusters in that condition. This analysis revealed several medium-specific enrichment patterns. For example, in TSB50, RiPP-like clusters were significantly enriched among expressed BGCs relative to other BGC types (FDR < 0.01), whereas terpene clusters showed strong activation in all media except SMM-gly. NI-siderophore clusters were significantly associated with DNPM and in all three defined media. T1PKS clusters were significantly enriched in most media except ISP2 and MA, while T3PKS clusters showed specific enrichment in SMM media with glycerol and maltose. In contrast, no significant association between media and expression was observed for other BGC types, including NRPS, NRPS-like, PKS-like, hglE-KS, ladderane, and lanthipeptide clusters. DNPM, TSB50, and SMM-malt triggered the most pronounced BGC-type–specific expression biases (Fig. 2D). These findings illustrate that while all the conditions tested support some level of specialized metabolism, in some cases, individual media clearly influence which types of BGCs become transcriptionally active.

### Role of cluster-specific regulators in BGC expression

We next examined whether specific gene cluster families (GCFs) exhibit consistent expression states (e.g. always expressed or consistently silent) across strains and assessed whether the presence or absence of cluster-specific regulators could explain these patterns. We grouped BGCs into GCFs using BiG-SCAPE v2 [36]. At a similarity threshold of 0.30, we identified 605 GCFs. For 145 (∼24%) of these GCFs, every BGC in the GCF was expressed in at least one media condition; for 185 (∼31%), only a subset of BGCs within a GCF were expressed; and for 275 (∼45%), every BGC within the GCF remained transcriptionally silent across all conditions (Fig. 3A, Supplementary Table 3). Increasing the clustering threshold to 0.40 or 0.50 increased the total number of GCFs (614 and 649, respectively), but the balance between fully expressed, partially expressed, and silent GCFs remained similar (Fig. 3A, Supplementary Tables 4 and 5, respectively). We adopted the more stringent 0.30 threshold for all subsequent references to GCFs in this study as it reduced the likelihood of unrelated BGCs being grouped into the same GCF. To investigate why some BGCs were expressed in only a subset of the strains that harbor them, we generated synteny plots for these GCFs using clinker (Fig. 3B–D) [37]. Specifically, we focused on GCFs defined at a 0.30 similarity threshold that contained at least four members and at least one regulatory gene within the BGC. To ensure high-quality clustering, we further required that core biosynthetic genes show consistent overlap in the synteny plots. In total, 41 GCFs met these criteria. Among these, we identified three cases in which differences in BGC expression correlated with variation in the presence or absence of cluster-associated regulatory genes.

**Fig. 3.**
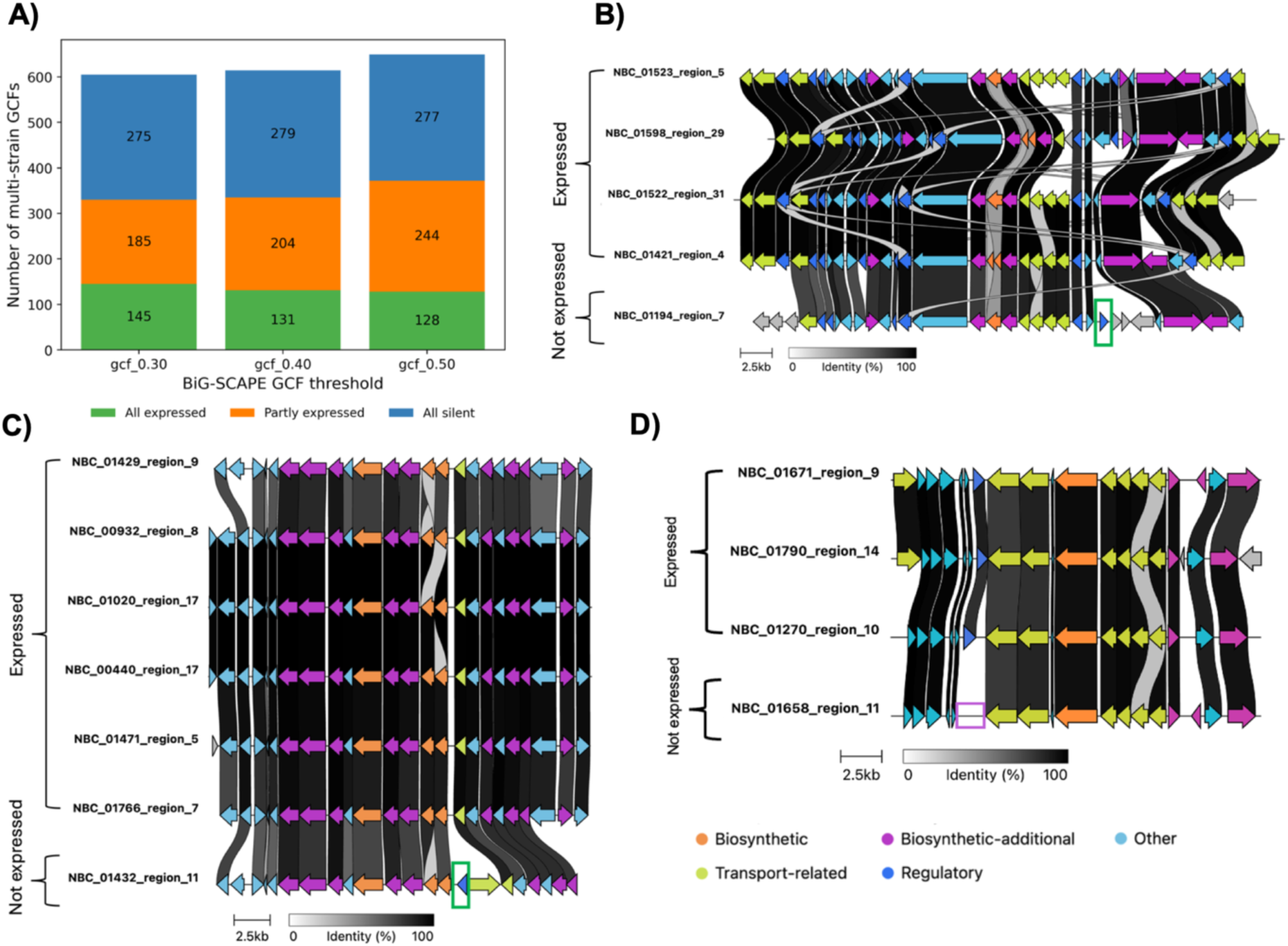
**A)** Number of GCFs found in at least two strains, across different similarity thresholds, categorized by whether all BGCs within a GCF are expressed, silent, or only partially expressed. **(B–D)** Representative examples of partially expressed BGCs within a GCF visualized as clinker synteny plots. **B)** GCF 1310: the silent cluster has additional TetR/AcrR family transcriptional regulator (green box). **C)** GCF 1274: the silent cluster has additional TetR/AcrR family transcriptional regulator (green box). **D)** GCF 1658: the silent cluster lacks a response regulator TF (purple box).

For example, in GCF 1310, which comprises T3PKS clusters, the silent cluster contains an additional TetR/AcrR-family transcriptional regulator that is absent in the expressed BGCs (Fig. 3B). A similar pattern appears in GCF 1274, comprising hopanoid BGCs: the non-expressed BGC harbors an extra TetR/AcrR-family regulator compared to expressed versions of the cluster (Fig. 3C). In contrast, GCF 1658, comprising SAL-242 lanthipeptide BGCs, shows the opposite trend: the silent cluster lacks a response regulator transcription factor (TF) that is present in the expressed BGCs (Fig. 3D). Together, these examples suggest that strain-specific gain or loss of cluster-associated regulatory genes can contribute to differential activation of conserved BGCs. In the remaining GCFs, no consistent association between BGC expression and the presence or absence of cluster-encoded regulatory genes was observed.

### Genes that are most commonly co-expressed with BGCs

We next investigated which genes were consistently co-expressed with BGCs by constructing gene co-expression networks using iterativeWGCNA [33], which is a variant of standard WGCNA [32]. Both methods group genes with highly correlated expression patterns into modules, but iterativeWGCNA refines the modules to improve robustness. Co-expression networks were built for each strain and spanned all media conditions, after which the data across all strains were aggregated and analyzed collectively. Furthermore, we used the RNA-seq data to define BGC borders more accurately prior to building these networks so that co-expression analysis would exclude genes not functionally part of the cluster. Across the 132 strains, there were 15,938 co-expression modules in total (median 109 modules per strain; IQR 90.8–137, range 28–296), with the majority of genes assigned to modules (median 3.48% unassigned per genome). These modules were then tested to identify those significantly enriched for BGC genes. We identified 895 BGC-enriched modules across the dataset, collectively associated with 1024 distinct BGCs (Supplementary Table 6).

We classified co-expressed genes based on their gene product annotation and orthogroup [38], and then counted the number of distinct BGCs with which they were co-expressed. At the gene product level (Fig. 4, Supplementary Table 7), the functions most frequently co-expressed with BGCs were consistent with known requirements for specialized metabolism. These included membrane transporters, particularly MFS and ABC transporter components, which may contribute to precursor uptake, metabolite export, or self-resistance [39–42]. Regulatory genes, including diverse TF families, response regulators, and sigma factors, were also frequently associated with BGC expression. Other recurrent functions included redox and tailoring enzymes, such as oxidoreductases, cytochrome P450s, and methyl-transferases. Interestingly, classical nutrient-responsive regulators were not prominent. Phosphate limitation is a well-established trigger of specialized metabolism mediated through the PhoR–PhoP two-component system [7, 8], which regulates the pstSCAB operon [43]. In our dataset, pstSCAB was co-expressed with just 22 BGCs. This suggests that phosphate-dependent activation is not a dominant or generalizable driver of BGC expression. Importantly, co-expression analysis links BGCs to specific candidate regulatory and transport genes, providing information not available from genome annotation alone.

**Fig. 4.**
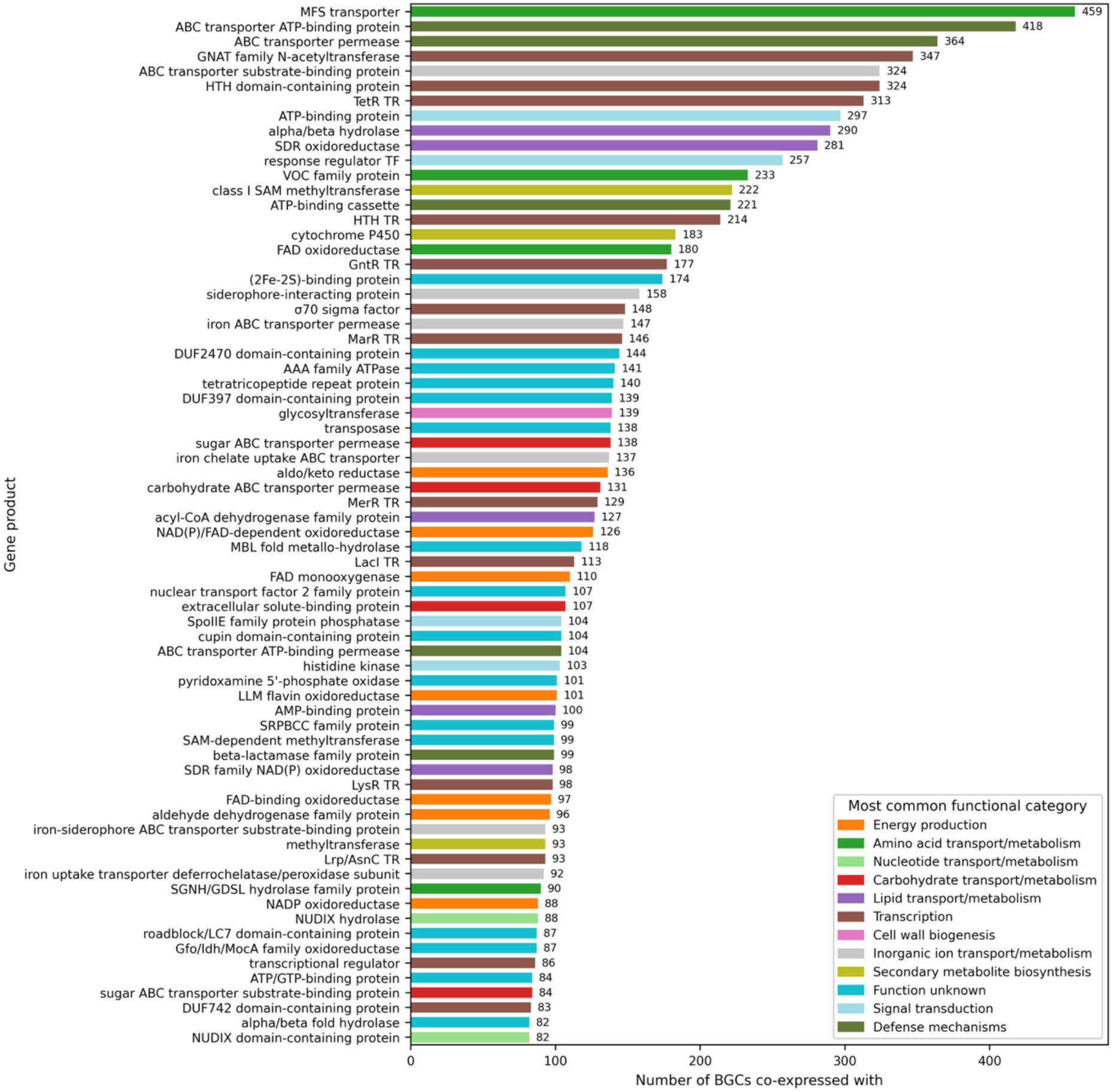
Top 70 gene products most frequently co-expressed with BGCs. Bars indicate the number of unique BGCs associated with each product annotation and are colored by the most common COG category among genes assigned to that product.

In addition to these well-characterized functions, several protein families of poorly characterized function were consistently co-expressed with BGCs. DUF2470-, DUF397-, and DUF742-containing proteins were co-expressed with 144, 139, and 83 BGCs, respectively (Fig. 4). DUF397-containing proteins were detected in 131 out of 132 strains, with a median of 26 per strain. DUF742-containing proteins were detected in 130 out of 132 strains, with a median of 10 per strain. Within BGC-associated modules, DUF397-containing genes and DUF742-containing genes were found in 83 and 59 strains, respectively (Supplementary Table 8). This suggests that they might act as overlooked regulators of BGC expression in diverse strains. Several other DUF containing proteins appeared less often but were associated with BGC expression at frequencies comparable to or exceeding that of phosphate-related regulation. For example, DUF4232-, DUF2637-, DUF485-, DUF4190-, and DUF5134-containing proteins were co-expressed with 44, 36, 35, 32, and 30 BGCs, respectively. Besides DUFs, 8663 hypothetical proteins were identified among BGC-associated module genes. Of these, 1970 had high module membership values (kME ≥ 0.95), indicating that their expression profiles closely followed the eigengene of their assigned module. These genes represented 1331 orthogroups, including 155 singletons, suggesting that BGC-associated modules contain many uncharacterized genes that may be specific to BGCs or strains (Supplementary Table 6).

When genes were grouped by orthogroup (Supplementary Fig. 3, Supplementary Table 9), conserved gene families corresponding to transport systems, transcriptional regulators, and redox or tailoring enzymes were repeatedly co-expressed with large numbers of BGCs. This consistency supports the robustness of the observed associations.

Among orthogroups enriched in genes annotated as hypothetical proteins, one orthogroup (OG0000027) stood out as being co-expressed with 53 distinct BGCs. Although most members of this orthogroup lack detailed annotation in GenBank, EggNOG-based predictions consistently suggest DNA-associated functions, including sequence-specific DNA binding and integrase- or recombinase-like activities. Several proteins also contain tetratricopeptide repeat domains. Notably, annotations of viral genome integration in bacterial genomes often arise from homology to integrase-like proteins and frequently correspond to cryptic or domesticated mobile elements rather than active prophages [44]. Tetratricopeptide repeat domains are well-established structural motifs that mediate protein–protein interactions in diverse regulatory complexes [45]. Taken together, these features suggest that OG0000027 represents a conserved regulatory or scaffolding module involved in higher-order transcriptional control rather than a biosynthetic enzyme family.

### Siderophore and metallophore associated BGCs have a distinct co-expression signature

We next examined whether certain BGC classes exhibited specific co-expression signatures, finding clear class-specific patterns in which distinct gene sets associated with specific cluster types rather than being broadly shared across specialized metabolism. These relationships were quantified using co-expression enrichment (log10 odds ratios; Fig. 5) and were repeated at the orthogroup level (Supplementary Fig. 4).

**Fig. 5.**
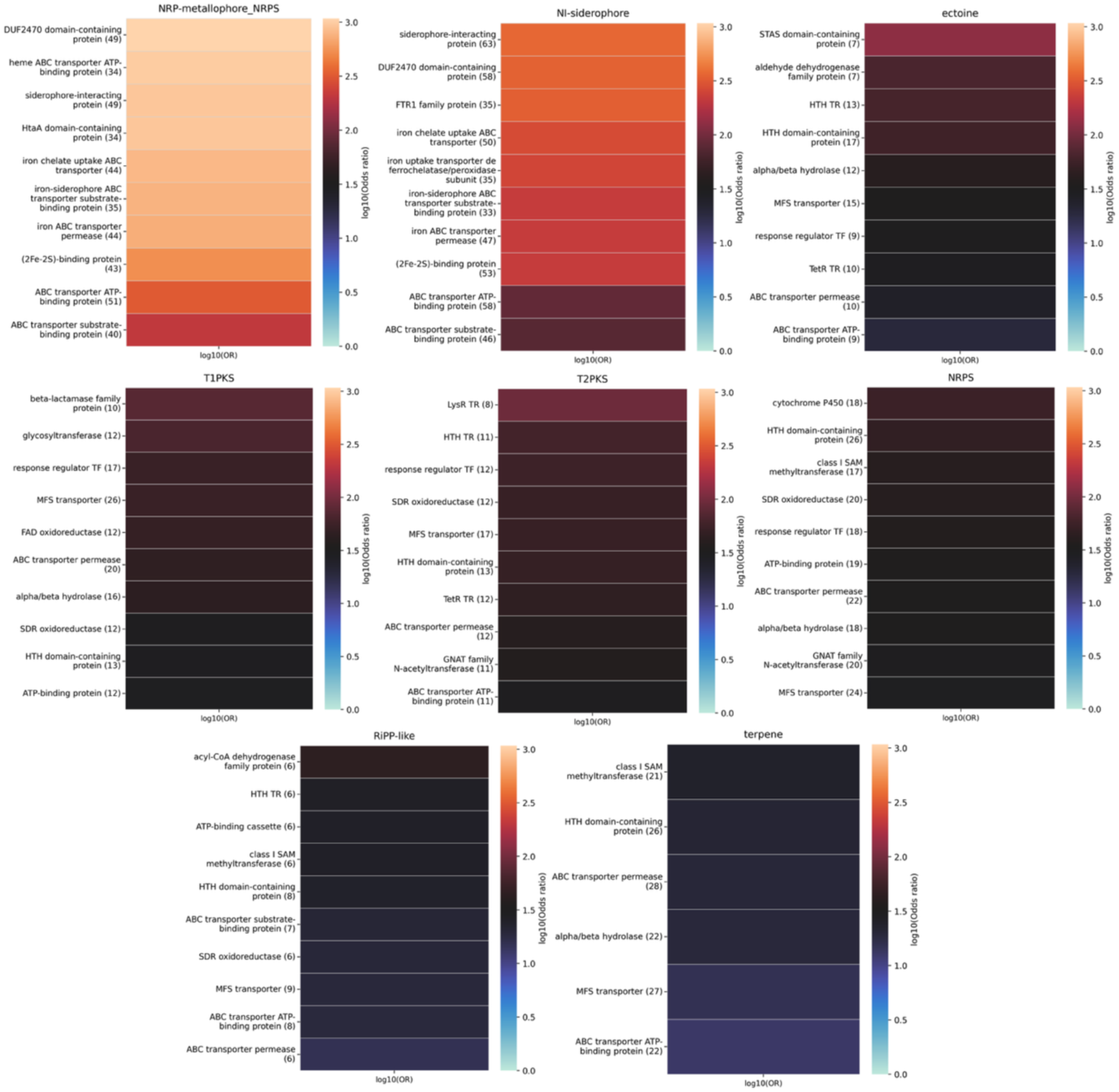
BGC-type–specific gene co-expression enrichment across BGC classes. Heatmaps show the top genes most strongly enriched in co-expression with each BGC type, quantified as log10 odds ratios. Rows represent individual genes, and colors indicate the strength of enrichment. Numbers shown in parentheses next to each gene product indicate the number of clusters of the corresponding BGC type with which that gene product is co-expressed. Only the top 10 enriched genes per BGC type are shown.

Both NRP-metallophore-NRPS (Fig. 5A) and NI-siderophore modules (Fig. 5B) were highly enriched for genes directly related to iron acquisition and handling. DUF2470 domain containing proteins appeared frequently and ranked among the top enriched functions for both classes. Others include (2Fe–2S)-binding proteins, iron-chelate uptake ABC transporters, iron ABC transporter permeases, siderophore-interacting proteins, and ferredoxin-associated redox components. This observation supports a tightly coordinated transcriptional program dedicated to metal acquisition and redox balance. In contrast, the remaining BGC types showed less coherent enrichment patterns in their modules. Although modules containing T2PKS, RiPP-like, NRPS, T1PKS, ectoine, and terpene clusters were enriched for some gene products, these signals were generally more heterogeneous and less specific than those observed for NI-siderophore and NRP-metallophore/NRPS-like clusters (Fig. 5C-H, Supplementary Table 10).

Analysis of enrichment at the orthogroup level mirrored these same trends for NI-siderophore and NRP-metallophore-NRPS clusters, demonstrating that this co-expression signature is conserved across homologous genes in different strains (Supplementary Fig. 4). In addition, two orthogroups dominated by proteins annotated as hypothetical were significantly correlated with NRP-metallophore-NRPS expression. The first, OG0005193 (co-expressed with 15 NRP-metallophore-NRPS BGCs, Supplementary Table 11), is annotated as hypothetical in GenBank; however, EggNOG-based functional prediction assigns this orthogroup a putative amine dehydrogenase–related activity, suggesting a possible enzymatic role that has not yet been experimentally validated. The second, OG0003886 (co-expressed with 11 NRP-metallophore-NRPS BGCs, Supplementary Table 11), lacks functional annotation in both GenBank and EggNOG. The recurrent enrichment of these poorly characterised orthogroups alongside metallophore-associated BGCs highlights conserved gene families that may contribute to metal acquisition or regulation but whose functions have not yet been characterized.

### Overexpression of genes co-expressed with BGCs enhances specialized metabolite output

We next tested whether genes that were co-expressed with BGCs, when overexpressed, could directly influence specialized metabolite production levels. All genes appearing within a co-expression module containing a BGC were candidates for overexpression, but we prioritized those with high correlation with the module eigengene (kME) and those with putative regulatory roles, including hypothetical proteins and genes annotated with transcription-related COG categories. Furthermore, we overexpressed the DUF397- and DUF742-containing operons from the granaticin-associated module in *Streptomyces sp.* NBC_00223 in all tested strains. Both were prioritized because DUF397 and DUF742 were co-expressed frequently with many BGCs across many strains, suggesting a potential broader role in specialized metabolism. The DUF397 operon comprised genes encoding a hypothetical protein and a DUF397-domain protein, whereas the DUF742 operon comprised genes encoding a nitrate/nitrite-sensing protein, a roadblock/LC7-domain protein, a DUF742-domain protein, and an ATP/GTP-binding protein. For all overexpression experiments, production levels were quantified relative to the empty-vector control unless otherwise stated. Media were selected based on BGC expression observed in the wild-type (WT) strain. Full locus identifiers, annotations, co-expression module names and kME can be found in Supplementary Table 12. Specific modules can be found in Supplementary Table 6.

We first targeted a co-expression module containing a BGC in *Streptomyces* sp. NBC_00906 associated with production of a red compound. The link between the BGC and the red pigment was confirmed by generating a knockout mutant of a core biosynthetic gene, which abolished pigment production (Supplementary Fig. 5, Supplementary Results and Methods). The module comprised 243 genes, including the complete BGC. In addition to the DUF742 and DUF397 operons, three additional candidates were selected for overexpression: a sigma/anti-sigma operon (Sig11105–AntiSig11110), a hypothetical protein (HP43505), and a GntR-family TF (GntR07510).

In DNPM medium (Fig. 6A), overexpression of the DUF742 operon resulted in a 3.0-fold increase in production of the red compound (P < 0.01). Overexpression of the DUF397 operon led to a stronger effect, with a 4.7-fold increase in production (P < 0.001). In the defined SMM-gly medium (Fig. 6B), baseline pigment production was lower than in DNPM. Overexpression of the DUF742 operon resulted in a 6.3-fold increase in red compound production (P < 0.01), while DUF397 operon overexpression led to a 5.0-fold increase (P < 0.01). Additionally, overexpression of the Sig11105–AntiSig11110 resulted in a 2.0-fold increase in pigment production (P < 0.01). Raw pellet weight and absorbance measurements are provided in Supplementary Table 13.

**Fig. 6.**
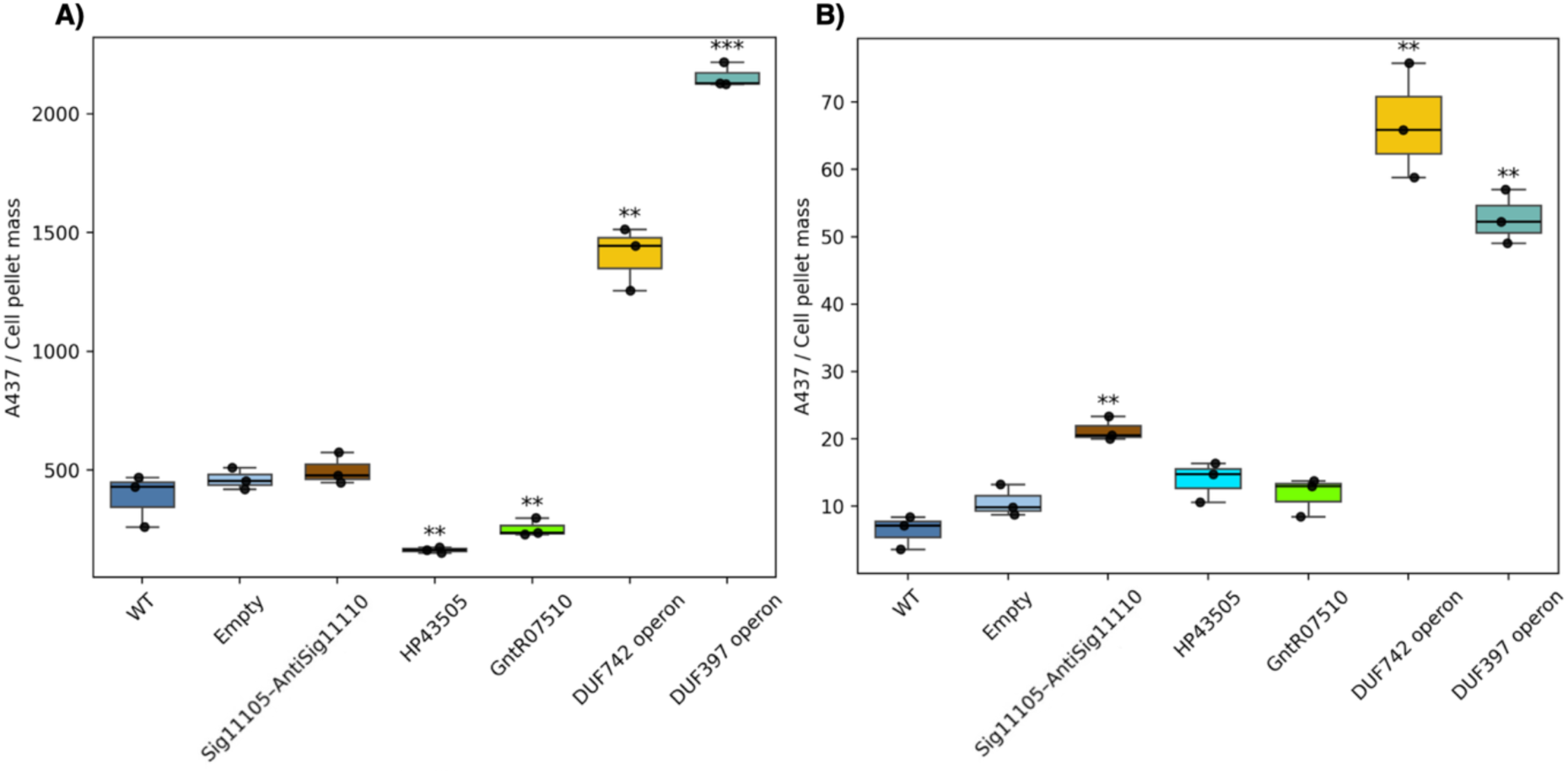
Effects of candidate gene and operon overexpression on red compound production. **A)** Red compound production in DNPM medium at 96 h. **B)** Red compound production in SMM–gly medium at 96 h. Red compound absorbance at 437 nm was normalized to cell pellet mass. Boxes show the interquartile range with the median indicated; points represent biological replicates. Asterisks indicate significant differences relative to empty vector (Student’s t-test; P < 0.05 *, P < 0.01 **, P < 0.001 ***; Benjamini–Hochberg FDR applied).

We next focused on two co-expression modules in *S. coelicolor* strain NBC_00116 associated with the actinorhodin (ACT) and undecylprodigiosin (RED) BGCs (Fig. 7). The ACT-associated module comprised 52 genes, including the complete ACT BGC, while the RED-associated module comprised 66 genes, including the complete RED BGC. From the ACT module, we overexpressed the DUF742 and DUF397 operons again along with a SigE family sigma factor (SigE18100) and a hypothetical protein (HP22130). From the RED module, we overexpressed the DUF742 and DUF397 operons, the operon containing AfsR, PadR, and a hypothetical protein (OP04285–04295) and another hypothetical protein by itself (HP22095). ACT and RED were both quantified over time because these metabolites are produced sequentially by *S. coelicolor*, with RED preceding ACT [46].

**Fig. 7.**
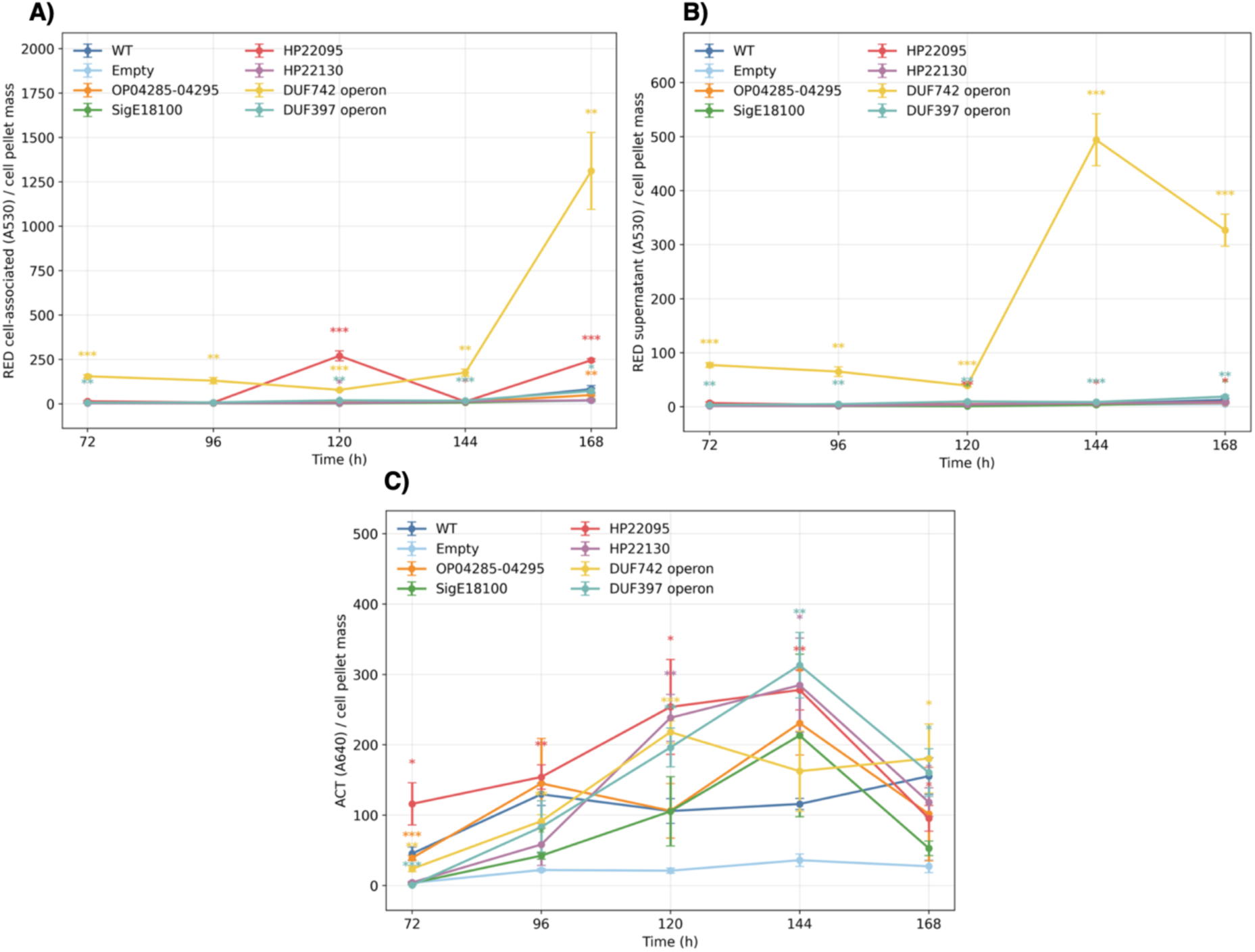
Effects of candidate gene and operon overexpression on RED and ACT production. RED production in DNPM medium measured in **A)** the cell pellet and **B)** supernatant. **C)** ACT production in ISP2 medium. Points show mean ± SEM. Asterisks indicate significant differences relative to the empty vector at the same time point (Student’s t-test; P < 0.05 *, P < 0.01 **, P < 0.001 ***; Benjamini–Hochberg FDR applied).

For RED, we measured metabolite levels in both the cell pellet and supernatant. In the cell pellet in DNPM medium, more pronounced effects were observed. DUF742 operon overexpression resulted in extremely large increases in cell-associated RED, including 41-fold at 120 h and 74-fold at 168 h (P < 0.01), with some induction evident at earlier time points (33–54-fold at 72–96 h) (Fig. 7A). DUF397 operon again showed significant increases, including a 10.3-fold increase at 120 h and 4.1-fold at 168 h (P < 0.05). HP22095 resulted in strong intracellular increases at later time points, reaching 141-fold at 120 h and 11.9-fold at 168 h (P < 0.01).

In the extracellular fraction in DNPM medium, DUF742 operon overexpression led to dramatic increases in RED levels across all time points, ranging from 12.6-fold at 120 h to over 114-fold at 144 h and 66.6-fold at 168 h (P < 0.01) (Fig. 7B). Even at early time points (72–96 h), DUF742 operon induced strong increases of 33–55-fold (P < 0.05). DUF397 operon also significantly enhanced extracellular RED production, albeit to a lesser extent, with consistent 2–3.8-fold increases across the time course (72–168 h; P < 0.05). Overexpression of HP22095 increased extracellular RED levels by 1.6-fold at 120 h and up to 1.8-fold at 168 h (P < 0.05), while SigE18100 displayed a significant reduction at 120 h (1.3-fold, P < 0.05) followed by a modest increase at 168 h (1.7-fold, P < 0.05).

For ACT, the empty-vector control exhibited reduced production compared to the WT strain in ISP2 medium, reflecting possible metabolic burden associated with plasmid carriage, so the overexpression strains were compared to both WT and empty control. Relative to the empty vector, overexpression of HP22095 at 72 h resulted in a 32-fold increase in ACT production (P < 0.05), while overexpression of DUF742 and OP04285-04295 operons resulted in 6.5-(P < 0.05) and 10.8-fold (P < 0.01) increases, respectively (Fig. 7C). At 96 h, DUF742, SigE18100, and HP22095 showed moderate increase in ACT production (2-fold P < 0.05,4.2-7.0-fold, P < 0.01). HP22130 showed a delayed increase in ACT production with no significant effect at 72 h or 96 h, but significant increases at 120 h (11-fold, P < 0.01), 144 h (8-fold, P < 0.05), and 168 h (4-fold, P < 0.05). The strongest and most consistent effects were observed at 120 h and 144 h. Compared to the WT strain at these time points, several overexpression constructs enhanced ACT production. DUF742 operon and HP22095 overexpression resulted in 2.1–2.4-fold increases at 120 h (with DUF742 operon reaching statistical significance: P < 0.01), while HP22095 and DUF397 operon showed 2.4–2.7-fold increases at 144 h (P < 0.01, P < 0.05). These results indicate that overexpression of selected module-associated genes can not only restore ACT production relative to the empty-vector control but also surpass WT production levels (Supplementary Tables 14 and 15).

As a final example, we examined a co-expression module associated with the granaticin BGC in *Streptomyces sp.* NBC_00223 (Fig. 8). This module comprised 140 genes, including the complete granaticin BGC. In addition to the DUF742 operon and DUF397 operon, three genes were selected: transcription termination factor Rho (Rho31155), hypothetical protein (HP12820), and Sigma70 family sigma factor (Sigma70-15275).

**Fig. 8.**
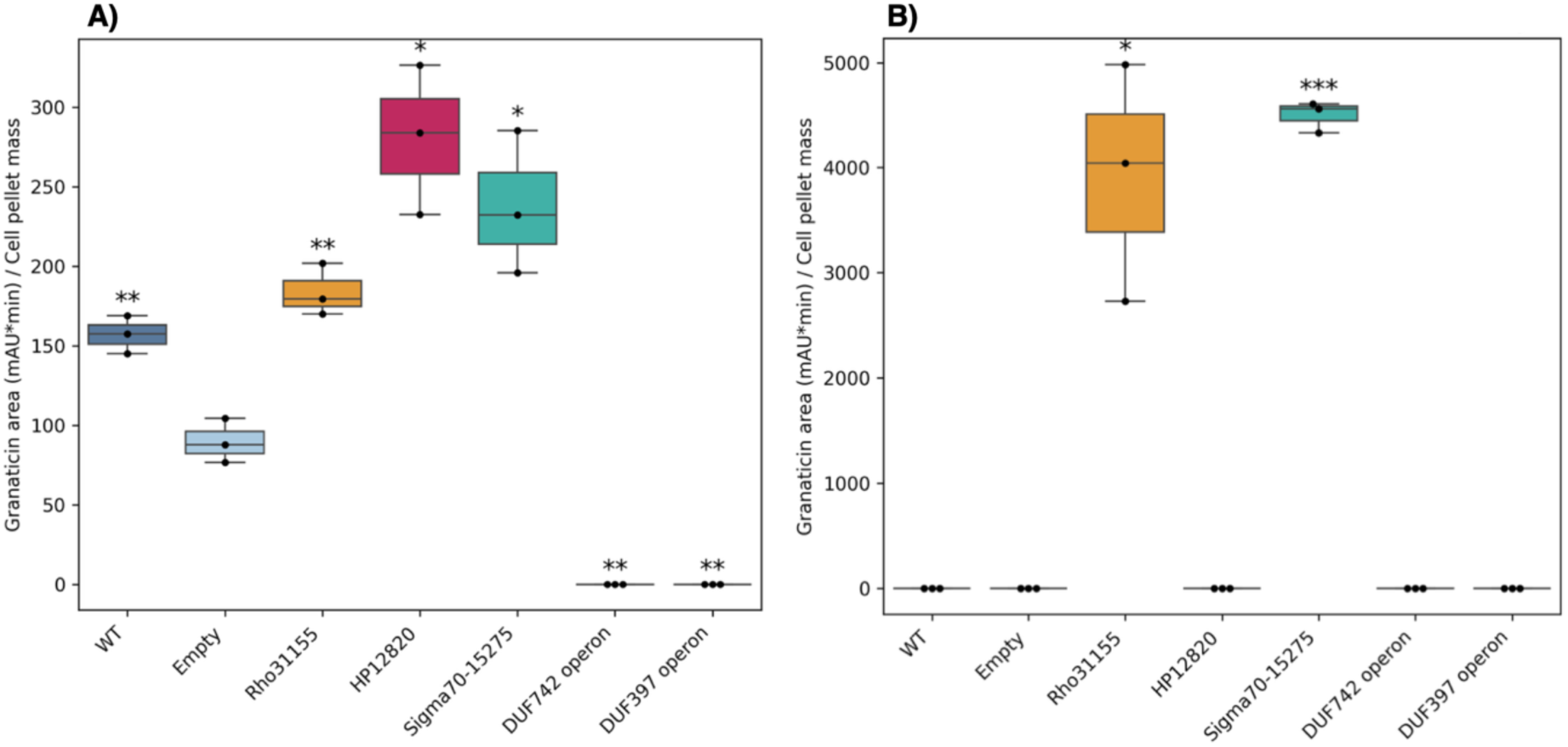
Effects of candidate gene and operon overexpression on granaticin production. **A)** Granaticin production in DNPM medium at 168 h. **B)** Granaticin production in SMM–gly medium at 288 h. Granaticin area (mAU·min) was normalized to cell pellet mass. Boxes show the interquartile range with the median indicated; points represent biological replicates. Asterisks indicate significant differences relative to WT (Student’s t-test; P < 0.05 *, P < 0.01 **, P < 0.001 ***; Benjamini–Hochberg FDR applied).

In DNPM medium (Fig. 8A), granaticin production levels were higher in the WT than empty vector control. Relative to the empty-vector control, overexpression of HP12820 resulted in a 3.1-fold increase in granaticin production (P =< 0.05), representing the strongest enhancement among the tested candidates. Overexpression of Sig70-15275 also increased production (2.7-fold, P < 0.05), while Rho31155 produced a more moderate effect (2.1-fold, P < 0.01). In contrast, overexpression of the DUF742 and DUF397 operons did not lead to detectable granaticin production under these conditions. In SMM-gly medium (Fig. 8B), neither the WT nor the empty-vector control produced detectable granaticin. However, overexpression of Rho31155 and Sig70-15275 led to robust granaticin production. The raw HPLC peak areas for granaticin and corresponding cell pellet measurements are provided in Supplementary Table 16.

Overall, increased metabolite production was observed for several overexpression candidates, with the DUF742 and DUF397 operons showing the broadest effects across multiple BGCs, while other candidates displayed more metabolite-specific responses (Table 1).

**Table 1.**
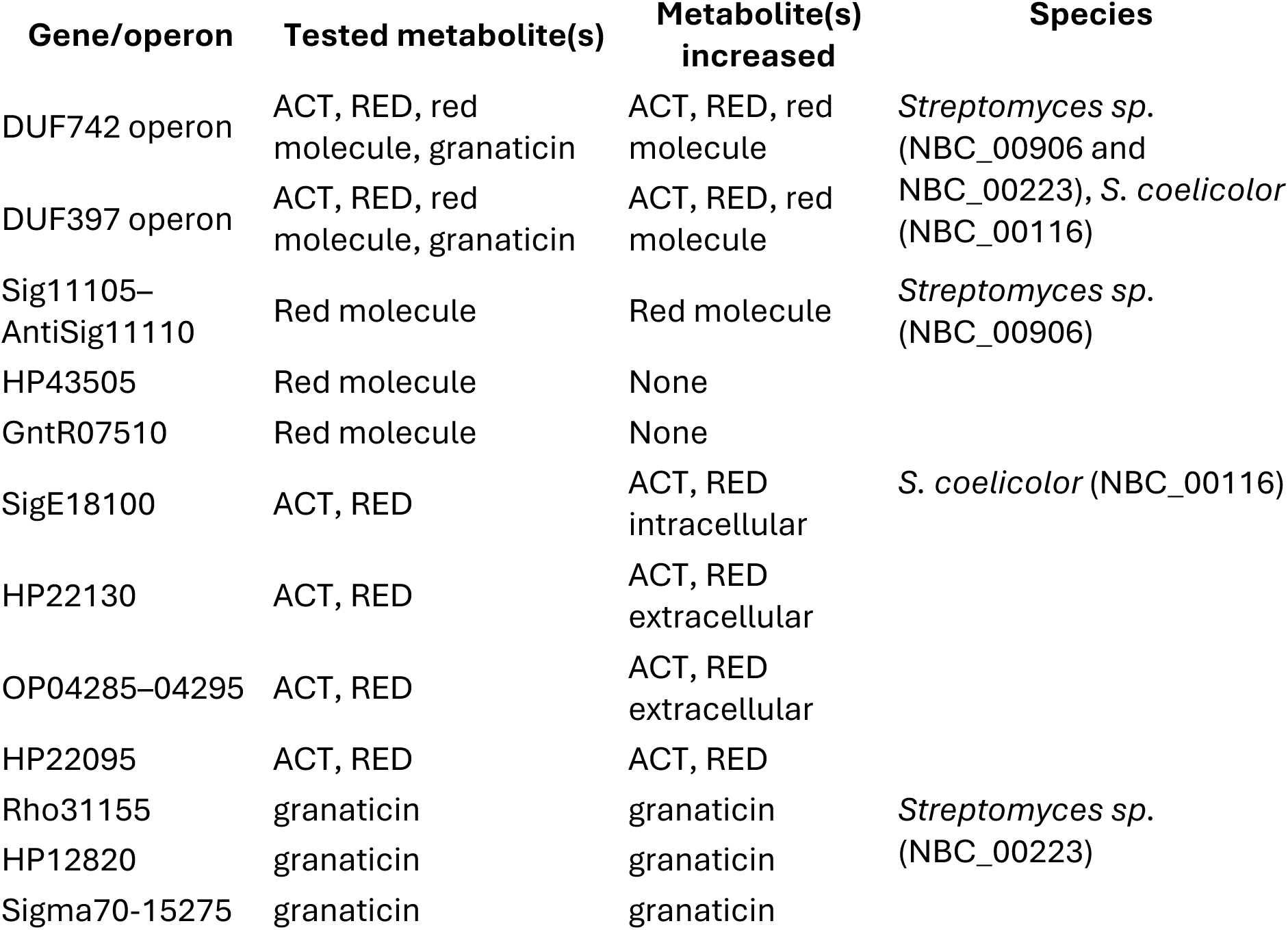
Summary of changes in metabolite levels following overexpression of genes co-expressed with four BGCs relative to the empty vector. Increases indicate higher production in at least one tested medium or time point.

## DISCUSSION

In this study, we generated an RNA-seq compendium encompassing 132 Actinomycetota strains and 1432 transcriptomic datasets covering different media. By integrating genome mining with co-expression analysis, we mapped BGC expression and identified genes co-expressed with BGCs. Overall, 43.7% of BGCs were expressed in at least one condition, with known BGCs more frequently expressed than unknown BGCs (61% versus 35%). Comparative analysis of GCFs suggested that strain-specific regulatory architecture can contribute to differential activation of BGCs. Co-expression analysis revealed conserved transcriptional patterns for siderophore and metallophore BGCs and frequent associations between BGCs and DUF742- or DUF397-containing genes. Experimental validation showed that several BGC-associated genes can increase specialized metabolite production, including genes annotated as hypothetical.

Although BGCs are often described as silent or cryptic, our data show that BGC expression is more widespread than is commonly assumed. Nearly half of all encoded BGCs were transcribed in at least one medium, consistent with previous work in *Salinispora* that reported a similar level of BGC expression [47]. However, expression was unevenly distributed: known BGCs were more frequently expressed than unknown BGCs. This pattern may partly reflect discovery bias, because BGCs with high expression, abundant associated metabolite production, or clear biological functions are more likely to have been detected and characterized previously. At the same time, our dataset provides experimentally supported expression conditions for over 1,000 previously uncharacterized BGCs, offering a direct route to prioritize clusters for specialized metabolite discovery. For many clusters, the remaining barrier to compound discovery may not be transcriptional activation alone, but may also involve translation, metabolite stability, compound abundance, extraction efficiency, or detection methods.

BGC expression varied across media, but was rarely restricted to a single growth condition, suggesting that activation often reflects overlapping physiological cues rather than highly specific medium triggers. At the GCF level, strain-specific activation coincided with differences in regulatory architecture, including the presence or absence of cluster-associated regulators. These observations suggest that BGC activation depends not only on the external growth condition, but also on the regulatory configuration of the BGC within each strain.

Co-expression analysis revealed that BGCs are co-expressed with transport-related genes more than any other gene category, and that the transporters are frequently located outside predicted BGC boundaries. Notably, transport-related functions comprised four of the top five co-expressed gene product categories (Fig. 4). The breadth of this association is consistent with transporter roles in precursor uptake, metabolite export, and self-resistance [39–42]. Co-expression analysis can thus link individual BGCs with distally encoded transporters that would be difficult to assign based on genome annotation alone, providing a practical route to identify specific transporters that may support metabolite production. Overexpression of such transporters could be a promising strategy to boost the yield of specialized metabolites during strain engineering.

Although DUF397- and DUF742-containing operons have previously been linked to specialized metabolism, our analysis extends these observations. DUF397 proteins are commonly encoded downstream of XRE-family transcription factors, forming conserved XRE-DUF397 regulatory pairs that are widespread in *Streptomyces* [13, 14]. DUF742 proteins are associated with conservon systems, membrane-associated signalling complexes implicated in development and specialized metabolism [15, 16]. DUF397-and DUF742-containing genes appeared in BGC-associated co-expression modules in 83 and 59 out of the 132 strains, respectively, suggesting that they have broader and more general roles in BGC regulation. However, there are often multiple copies of each in a genome, making it difficult to pinpoint the precise copy that is associated with a particular BGC based on annotation alone. Co-expression analysis, therefore, provides a way to prioritize the specific DUF397- or DUF742-containing operons most likely to influence BGC expression or metabolite production. Additionally, we identified many hypothetical genes and orthogroups co-expressed with BGCs, suggesting that BGC-associated modules contain numerous uncharacterized genes that may contribute to specialized metabolism as well.

NI-siderophore and NRP-metallophore/NRPS-like clusters were repeatedly co-expressed with similar sets of genes, including siderophore-interacting proteins, iron uptake systems, Fe--S-binding proteins, redox enzymes, and DUF2470-domain proteins. This coherent signature likely reflects the shared biological function of these clusters in metal acquisition, which requires coordinated regulation of biosynthesis, uptake, transport, and iron-responsive metabolism. Recent studies link DUF2470-family proteins to heme binding and iron homeostasis [48, 49]. Together, these observations suggest that DUF2470 proteins may participate in the broader regulatory or physiological context of siderophore and metallophore biosynthesis.

Collectively, our results support a multilayered model of BGC regulation in Actinomycetota in which activation is shaped by (i) environmental inputs such as medium composition, (ii) strain-specific regulatory genes withing BGCs in conserved GCFs, and (iii) integration of BGCs into broader transcriptional network modules. The substantial pool of conserved yet transcriptionally silent clusters underscores that regulatory architecture, rather than biosynthetic capacity alone, limits metabolite expression under laboratory conditions. By combining large-scale transcriptomics with co-expression network analysis and targeted perturbation, this study provides both a conceptual framework and a reusable resource for rational activation, prioritization, and engineering of specialized metabolism in *Streptomyces* and other Actinomycetota genera.

## METHODS

### Cultivation media and growth conditions

Most strains (n=117) were cultured in eight media as medium composition is known to influence specialized metabolite production [50]. These eight media included five complex media (DNPM [51], ISP2, MA [52], SoyM50 and TSB50) and three defined Streptomyces minimal media [53] supplemented with glucose (SMM-gluc), glycerol (SMM-gly) or maltose (SMM-malt) as the sole carbon source. The complex media were included to represent cultivation conditions commonly used for Actinomycetota growth and metabolite screening, whereas the defined SMM media allowed assessment of carbon source dependent changes in BGC expression under otherwise controlled nutritional conditions [54]. SoyM50 and TSB50 were prepared at half of their standard concentrations to test whether reduced nutrient availability could promote BGC expression. A subset was profiled across seven (n = 8) or fourteen (n = 7) conditions. The latter included two additional rich media (NB and AF [55]) and five additional defined media (SMM-ribose, SMM-mannose, SMM-starch, SMM-avicel, and SMM-xylose). Among these seven strains, six were sampled in SMM-ribose, SMM-mannose, SMM-starch, SMM-avicel, and SMM-xylose, whereas one strain was sampled in SMM-starch instead of SMM-avicel. Detailed media compositions are provided in Supplementary Table 17.

All strains were initially grown as pre-cultures in 50 mL ISP2 medium in 300 mL baffled flasks. Pre-cultures were incubated at 30°C with shaking at 140 rpm until sufficient biomass was obtained. For experimental cultures, 30 μL of the pre-culture was used to inoculate 30 mL medium in 100 mL Erlenmeyer flasks sealed with cotton plugs and aluminum foil. Cultures were subsequently incubated again at 30°C with shaking at 140 rpm. The number of days each strain was grown prior to RNA extraction is provided in Supplementary Table 18.

### RNA extraction and sequencing

After cultivation, we centrifuged the samples at 5,000 rpm for 4 minutes. The cell pellets were resuspended in 600 μL Buffer RLT supplemented with 2-mercaptoethanol (10 μL/mL), after which the suspensions were transferred to 2 mL Eppendorf tubes containing glass beads (Sigma Aldrich, 0.5 mm) filled to the 0.5 mL mark and disrupted using a Qiagen TissueLyser at 50 Hz for 5 minutes. After centrifuging the samples at 13,000 rpm for 1 minute to pellet the beads and all solid material, 350 μL of the lysate was transferred to a new 2 mL centrifuge tube. Special care was taken to avoid touching the glass beads and any solid material that lay on top with the pipet tip. Total RNA was then purified using the RNeasy Mini Kit (Qiagen) on the QIAcube automated platform using the “RNeasy Mini Animal Cells” protocol. A 15 minute on-column DNase I treatment was included as part of the purification. Buffers (70% ethanol, RW1, RPE, RNase-free water) and the DNase I/RDD solution were added according to the QIAcube’s guided setup. Purified RNA was kept at −80^◦^C for long-term storage.

RNA concentration and integrity were measured using Qubit 2.0 (Invitrogen) and Fragment analyzer (Agilent 530V). rRNA purification, library preparation and RNA sequencing were performed by Azenta Life Sciences (Leipzig, Germany). NEBNext rRNA Depletion kit (Bacteria) (New England Biolabs; cat. No. E7850X) was used for rRNA depletion. Barcoded libraries were prepared for each condition in duplicate with NEBNext Ultra II Directional RNA Library Prep Kit for Illumina using default protocols. Libraries for all samples were sequenced using Illumina NovaSeq 6000. An overview of all RNA-Seq samples, including strain, growth media, biological replicates, and mapping statistics, is provided in Supplementary Table 18.

### RNA-seq data processing and dataset quality

Raw RNA-seq reads were mapped to the reference genomes using a software package developed as part of independent component analysis [24]. This package consists of cutadapt to remove adapters low-quality sequences [56], Bowtie 1 for short read alignment to reference genomes [57], FastQC for quality control [58], RSeQC to evaluate additional quality metrics [59], and featureCounts for counting mapped reads [60]. Mapping statistics were summarised with MultiQC [61].

The initial dataset comprised 1568 RNA-seq samples from 137 Actinomycetota strains. Samples with fewer than one million reads assigned to features were excluded, resulting in a final compendium of 1432 samples from 132 strains. The strains span 12 genera, dominated by *Streptomyces* (109 strains), with additional representation from *Kitasatospora* (6), *Amycolatopsis* (4), *Micromonospora* (3), *Nocardia* (3), and seven other genera represented by one strain each (*Actinomadura*, *Embleya*, *Kribbella*, *Lentzea*, *Rhodococcus*, *Spirillospora*, *Streptomycetaceae*). To visualize the phylogenetic diversity of the strains, a genome-based phylogenetic tree was generated from GenBank files using getphylo [62] with default parameters and midpoint-rooted for visualization (Supplementary Fig. 6). As described, most strains were grown in eight different media. Most samples were represented by a single biological replicate, with additional biological replicates in selected conditions. Replicate consistency was assessed using Pearson correlation of log-TPM expression profiles, showing high concordance (mean r = 0.93 ± 0.06, median r = 0.95) (Supplementary Table 19). Sequencing yielded a median of 24.8 million reads per sample (IQR: 20.4–31.4 million), with a median of 10.4 million reads assigned to features. RNA integrity was high (median RQN = 9.5), and overall read quality averaged 35.8 ± 1.12. Across strains, a median of 99.3% of annotated genes had logTPM > 1 in at least one condition, indicating near-complete transcriptome coverage.

### Genome mining, functional annotation, and orthogroup inference

BGCs were identified using antiSMASH version 7.0 [34] and grouped into GCFs using BiG-SCAPE version 2.0 [36] based on domain composition and sequence similarity. At the gene level, protein-coding sequences were functionally annotated using eggNOG-mapper version 2 [63] based on EggNOG database version 5 [64]. Functional assignments included COG categories [65], providing broad biological classifications. To identify homologous gene families conserved across strains, orthogroups were inferred using OrthoFinder [38].

### BGC expression threshold

For each transcriptomic sample, we defined an expression threshold as the 75^th^ percentile of genome-wide expression values. A BGC was considered expressed if more than 50% of its core biosynthetic genes exceeded this threshold. Restricting the analysis to core genes reduces the impact of uncertain antiSMASH boundary predictions.

### Statistical enrichment analysis of BGC activation across media

Fisher’s exact tests were performed for each medium–BGC type combination. For every medium, a 2 × 2 contingency table was constructed. Specifically, *a* represented the number of expressed BGCs of the tested type, *b* the number of non-expressed BGCs of that type, *c* the number of expressed BGCs of all other types, and *d* the number of non-expressed BGCs of all other types within the same medium.

### IterativeWGCNA network construction and selection of growth conditions

Co-expression networks were constructed separately for each strain using the iterativeWGCNA R package [33]. To ensure comparability across strains, we first evaluated the impact of using different numbers of growth conditions for network inference. For the seven strains profiled across both 8 and 14 conditions, networks inferred from 14 conditions showed increased complexity, including more modules, a larger fraction of unassigned genes, and substantial reassignment of genes between modules. These results indicate that uneven sampling across media alters network structure and reduces comparability (Supplementary Table 20). Based on this analysis, we selected a standardized set of eight growth conditions for downstream network construction, while retaining strains with seven available conditions.

For each strain, an appropriate soft-thresholding power was estimated from the expression data using the findModules function in the PyWGCNA Python package, which applies standard expression filtering and selects a power based on the scale-free topology criterion [66]. The selected power was then used for iterativeWGCNA network construction.

All networks were built as signed networks, such that only positive correlations contributed to adjacency and module assignment. The minimum module size was set to 5 genes, and modules with eigengene correlation greater than 0.8 (cut-off = 0.2) were merged during iterative refinement. To ensure that all genes from a given strain could be analyzed in a single pass, the maxBlockSize parameter was set to approximately the number of genes in each genome. Full parameter settings and summary statistics for each strain-specific network are provided in Supplementary Table 21.

### Refinement of BGC boundaries using RNA-seq data

Within each predicted antiSMASH region, biosynthetic core genes were first grouped into core seed blocks based on genomic proximity. For each block, a mean expression profile (hereafter referred to as the BGC eigengene) was calculated to represent the characteristic transcriptional behavior of the biosynthetic core. Genes located within the same antiSMASH region were subsequently evaluated for inclusion in the refined BGC according to two criteria: (i) detectable expression (logTPM > 1 in at least one condition) and (ii) coordinated regulation with the biosynthetic core (Pearson correlation ≥ 0.6). This strategy enabled the identification of accessory, tailoring, and regulatory genes that are transcriptionally coupled to the core biosynthetic machinery. To preserve biologically meaningful cluster architecture, extension of cluster boundaries was constrained to maintain local genomic continuity, and small gaps between co-expressed genes were bridged.

### Definition of BGC-related co-expression modules

For each strain, each BGC, and each co-expression module, a 2 × 2 contingency table was constructed and Fisher’s exact test was performed. In this table, *a* represented the number of genes belonging to the BGC and assigned to the module, *b* the number of genes belonging to the BGC but not assigned to the module, *c* the number of genes not belonging to the BGC but assigned to the module, and d the number of genes neither belonging to the BGC nor assigned to the module.

Statistical significance alone is insufficient to distinguish biologically meaningful BGC-associated modules from large modules representing global transcriptional networks. Therefore, BGC-specific modules were defined using a combination of statistical, coverage and specificity criteria.

For each significant BGC–module association, two additional metrics were calculated. BGC coverage was defined as the fraction of BGC genes assigned to the module:

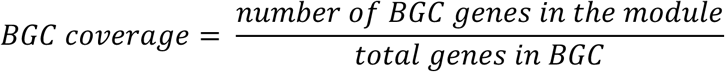

Module purity was defined as the fraction of module genes belonging to the BGC:

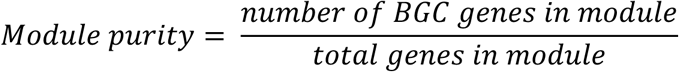

A co-expression module was classified as BGC-related if it satisfied all of the following criteria: (i) FDR < 0.01, (ii) BGC coverage ≥ 0.30, (iii) module purity ≥ 0.03, and (iv) at least three BGC genes assigned to the module. These were used to exclude large modules dominated by global physiological processes.

### Co-expression enrichment analysis for BGC types

For each gene–BGC type pair, a 2 × 2 contingency table was constructed and Fisher’s exact test was performed. In this table, *a* represented the number of observations in which the gene and the BGC type were co-expressed within the same module, *b* the number of observations in which the gene was present but not co-expressed with that BGC type, *c* the number of observations in which the BGC type was present without the gene, and d the number of observations in which neither was observed together.

For comparability across genes and BGC types, odds ratios were log_10_-transformed. To prevent extreme values driven by low counts from disproportionately influencing ranking and visualization, log-transformed values were capped at an upper bound. For each BGC type, genes were ranked according to their log_10_ odds ratio, and the top enriched genes were retained for downstream visualization. The same enrichment framework was independently applied at the orthogroup level.

### Construction of overexpression strains

For each gene that was overexpressed, primers were designed in StreptoCAD [67] to amplify the full-length coding sequence with overlaps compatible with the integrative overexpression vector pOEX-PkasO. In this plasmid, inserts are placed downstream of the strong constitutive *kasO\** promoter (P*_kasO_*_∗_) and an associated ribosome binding site, forming a defined expression cassette for high-level transcription in *Streptomyces*. PCR products were assembled into pOEX-PkasO following the standard cloning protocol implemented in StreptoCAD [67]. The resulting constructs were maintained in *E. coli* ET12567 donor cells and transferred into the corresponding host strains by intergeneric conjugation. After mating, exconjugants were selected on apramycin-containing media and purified by restreaking. Chromosomal integration of the plasmid was confirmed by diagnostic PCR. Correct assembly of the overexpression cassette was verified by Sanger sequencing. A complete list of overexpression constructs, amplified genes or operons, primer sequences is provided in Supplementary Table 12.

### Measurement of red compound, actinorhodin, undecylprodigiosin and granaticin

The red compound produced by the target BGC in strain NBC_00906 has UV absorbance at λ_max_ = 220 nm, 280 nm, 437 nm, and 550 nm. We quantified the red compound by measuring absorbance of the samples at 437 nm.

Quantification of ACT and RED in *S. coelicolor* cultures followed established spectrophotometric protocols with minor modifications [53]. For ACT, 1 mL of well-mixed culture was transferred to a microcentrifuge tube and supplemented with 250 μL of 5 M KOH. Samples were incubated overnight at 4 ^◦^C to convert actinorhodin to its blue, alkali-soluble form, then centrifuged at 13,000 ×g for 5 min. The absorbance of the supernatant was measured at 640 nm using a spectrophotometer, and the resulting values were used as a proxy for ACT production. For RED, cell pellets from 1 mL culture aliquots were collected by centrifugation and washed twice with 0.5 M HCl to remove extracellular pigment. The washed pellets were then extracted in 1 mL of methanol–HCl (0.5 M) and incubated for 2 h at room temperature. After centrifugation to remove cell debris, the absorbance of the supernatant was recorded at 530 nm as a proxy for RED production.

Granaticin production was quantified by high-performance liquid chromatography (HPLC) using a Thermo Ultimate 3000 system. Cultures were centrifuged and filtered, after which the supernatants were injected into a Waters XBridge C18 column (150 x 4.6mm, 3.5 μm pore size). Solvent A was water with 0.1% formic acid and 10 mM ammonium formate. Solvent B was 90:10 acetonitrile:water also with 0.1% formic acid and 10 mM ammonium formate. The mobile phase program was: 0-1 min isocratic 10% B; 1-11 min gradient from 10% to 100% B; 11-13 min isocratic 100% B; 13-14 min gradient from 100% to 10% B. Under these conditions, granaticin eluted at 7.9 minutes. Using wavelength 210 nm, the peak area at this retention time was automatically integrated and served as the measure of granaticin production.

To account for differences in metabolite levels stemming from differences in biomass accumulation, production measurements for each of the three compounds were normalized to dry cell mass for all strains and conditions. 2 mL of each culture were harvested, centrifuged at 5,000 rpm for 4 minutes to obtain cell pellets, and dried at 60^◦^C for 48 h. The resulting dry weight was used to normalize spectrophotometric and HPLC-based quantifications. This normalization was performed to ensure that differences in metabolite levels reflected changes in biosynthetic activity rather than variations in cell biomass.

### Statistical analysis

All statistical analyses were performed in Python. Enrichment analyses were conducted using Fisher’s exact tests implemented in the scipy.stats.fisher exact function of the SciPy library [68]. P-values were adjusted for multiple testing using the Benjamini–Hochberg procedure to control the false discovery rate (FDR) [69]. Associations with FDR < 0.05 were considered statistically significant.

## ACKNOWLEDGEMENTS

We thank Bernhard Palsson immensely for his support on this project. We also thank Chris Whitford for helpful advice on strain manipulation and overexpression experiments.

## FUNDING STATEMENT

A.R., T.W. and P.C. disclose support for the research of this work from Novo Nordisk Foundation through grants NNF20CC0035580 and NNF25SA0109652. H-S.K. discloses support of this work from the National Research Foundation of Korea (RS-2025-25460810). H.U.K discloses support of this work from the Bio & Medical Technology Development Program of the National Research Foundation unded by the Korean government (MSIT) (RS-2024-00352229 and RS-2025-02216377). The remaining authors declare no relevant funding.

## AUTHOR CONTRIBUTIONS

P.C. and A.R. initiated the study and extracted RNA samples. P.C. and O.M. mapped the RNA-seq data and did antiSMASH. A.R. designed and performed the computational analyses, including BiG-SCAPE, BGC expression profiling, GCF-level comparisons, co-expression network construction, and analysis of BGC-associated modules. B.T.L., B.L., and J.Y.K. contributed to the interpretation of computational analysis results. A.R. designed and carried out the experimental validation work, including candidate selection, overexpression experiments, and metabolite assays, with input from P.C. B.Y. helped with overexpression expreriments with *S. coelicolor* strain. P.C. collected HPLC data. H.U.K., A.S., H-S.K, and T.W. contributed to biological interpretation and manuscript revision. P.C. led the overall project. All authors interpreted the results, reviewed the manuscript, and approved the final version.

## COMPETING INTERESTS

The authors declare no competing interests.

## DATA AVAILABILITY

The genome sequences analysed during the current study are publicly available in the NCBI GenBank and RefSeq repositories [31]. Accession numbers for all genomes used in this study are provided in Supplementary Table 18. The RNA-seq data generated in this study have been deposited in the NCBI Sequence Read Archive (SRA) under accession number PRJNA1309148.

## CODE AVAILABILITY

The code used for BGC border refinement is publicly available on GitHub [https://github.com/arudenko-2025/BGC-refirement-RNA-Seq] and an archived on Zenodo [https://doi.org/10.5281/zenodo.18769627]. The custom code used for generation of figures and analysis for this manuscript is available at GitHub link [https://github.com/arudenko-2025/RNA-Seq-actinomycetota-data-analysis] and the version of code is also deposited at Zenodo [https://doi.org/10.5281/zenodo.20282452].

## Notes

### Competing Interest Statement

The authors have declared no competing interest.

## REFERENCES

[1] Baltz RH. Gifted microbes for genome mining and natural product discovery. J. Ind. Microbiol. Biotechnol. 44, 573–588 (2017). 10.1007/S10295-016-1815-X.

[2] van der Meij A, Worsley SF, Hutchings MI, van Wezel GP. Chemical ecology of antibiotic production by actinomycetes. FEMS Microbiol. Rev. 41, 392–416 (2017). 10.1093/FEMSRE/FUX005.

[3] Walsh CT, Fischbach MA. Natural Products Version 2.0: Connecting Genes to Molecules. J. Am. Chem. Soc. 132, 2469–2493 (2010). 10.1021/JA909118A.

[4] van Der Heul HU, Bilyk BL, McDowall KJ, Seipke RF, Van Wezel GP. Regulation of antibiotic production in Actinobacteria: new perspectives from the post-genomic era. Nat. Prod. Rep. 35, 575–604 (2018). 10.1039/C8NP00012C.

[5] Takano E, Gramajo HC, Strauch E, Andres N, White J, Bibb MJ. Transcriptional regulation of the redD transcriptional activator gene accounts for growth-phase-dependent production of the antibiotic undecylprodigiosin in *Streptomyces coelicolor* A3(2). Mol. Microbiol. 6, 2797–2804 (1992). 10.1111/j.1365-2958.1992.tb01459.x.

[6] Arias P, Fernández-Moreno MA, Malpartida F. Characterization of the pathway-specific positive transcriptional regulator for actinorhodin biosynthesis *in Streptomyces coelicolor* A3(2) as a DNA-binding protein. J. Bacteriol. 181, 6958–6968 (1999). 10.1128/jb.181.22.6958-6968.1999.

[7] Martín JF. Phosphate control of the biosynthesis of antibiotics and other secondary metabolites is mediated by the PhoR-PhoP system: an unfinished story. J. Bacteriol. 186, 5197–5201 (2004). 10.1128/JB.186.16.5197-5201.2004.

[8] Martín JF, Rodríguez-García A, Liras P. The master regulator PhoP coordinates phosphate and nitrogen metabolism, respiration, cell differentiation and antibiotic biosynthesis: comparison in *Streptomyces coelicolor* and *Streptomyces avermitilis*. J. Antibiot. 70, 534–541 (2017). 10.1038/ja.2017.19.

[9] Rigali S, Titgemeyer F, Barends S, Mulder S, Thomae AW, Hopwood DA, et al. Feast or famine: the global regulator DasR links nutrient stress to antibiotic production by *Streptomyces*. EMBO Rep. 9, 670–675 (2008). 10.1038/embor.2008.83.

[10] Chater KF, Chandra G. The evolution of development in *Streptomyces* analysed by genome comparisons. FEMS Microbiol. Rev. 30, 651–672 (2006). 10.1111/j.1574-6976.2006.00033.x.

[11] Uguru GC, Stephens KE, Stead JA, Towle JE, Baumberg S, McDowall KJ. Transcriptional activation of the pathway-specific regulator of the actinorhodin biosynthetic genes in *Streptomyces coelicolor*. Mol. Microbiol. 58, 131–150 (2005). 10.1111/j.1365-2958.2005.04817.x.

[12] Hoskisson PA, Rigali S, Fowler K, Findlay KC, Buttner MJ. DevA, a GntR-like transcriptional regulator required for development in *Streptomyces coelicolor*. J. Bacteriol. 188, 5014–5023 (2006). 10.1128/JB.00307-06.

[13] Santamaría RI, Sevillano L, Martín J, Genilloud O, González I, Díaz M. The XRE-DUF397 protein pair, Scr1 and Scr2, acts as a strong positive regulator of antibiotic production in *Streptomyces*. Front. Microbiol. 9, 2791 (2018). 10.3389/fmicb.2018.02791

[14] Riascos C, Fernández-García G, García-Martín J, Manteca Santamaría RI, Díaz M. Overexpression of Scr1 protein induces strong changes in the transcriptome and metabolome of Streptomyces coelicolor that affect antibiotic production. Microb. Cell Fact. 24, 218 (2025). 10.1186/S12934-025-02843-5.

[15] Bonet, B. et al. The cvn8 conservon system is a global regulator of specialized metabolism in *Streptomyces coelicolor* during interspecies interactions. mSystems 6, e00281–21 (2021). 10.1128/msystems.00281-21

[16] Morin, L. M. C., Dekoninck, K., Min, K. Y. & Traxler, M. F. The accessory protein CvnF8 modulates histidine kinase activity in an Actinobacterial G protein system in *Streptomyces coelicolor*. bioRxiv (2025). 10.1101/2025.07.03.663114

[17] Lee, B. T. et al. Pan-reactome analysis of *Streptomyces* strains reveals association and disconnection between primary and secondary metabolism. Metab. Eng. 92, 241–251 (2025). 10.1016/j.ymben.2025.08.005

[18] Mohite, O. S. et al. Pangenome mining of the *Streptomyces* genus redefines species’ biosynthetic potential. Genome Biol. 26, 9 (2025). 10.1186/s13059-024-03471-9

[19] Morehouse, N. J. et al. Annotation of natural product compound families using molecular networking topology and structural similarity fingerprinting. Nat. Commun. 14, 308 (2023). 10.1038/s41467-022-35734-z

[20] Linington, R. G. An assessment of chemical diversity in microbial natural products. ACS Cent. Sci. 11, 1536–1545 (2025). 10.1021/acscentsci.5c00804

[21] Augustijn, H. E., Roseboom, A. M., Medema, M. H. & van Wezel, G. P. Harnessing regulatory networks in Actinobacteria for natural product discovery. J. Ind. Microbiol. Biotechnol. 51, (2024). 10.1093/jimb/kuae011

[22] Wilbanks, L. E. et al. DAP-seq reveals cluster-situated regulator control of numerous *Streptomyces* natural product biosynthetic genes. J. Nat. Prod. 89, 1440–1453 (2026). 10.1021/acs.jnatprod.5c01605

[23] Augustijn, H. E. et al. A global regulatory atlas of *Streptomyces* reveals conserved and diversified transcriptional networks across actinomycetes. bioRxiv (2026). 10.64898/2026.06.06.730586

[24] Sastry, A. V. et al. The *Escherichia coli* transcriptome mostly consists of independently regulated modules. Nat. Commun. 10, 5536 (2019). 10.1038/s41467-019-13483-w.

[25] Lee, Y. et al. Machine-learning analysis of *Streptomyces coelicolor* transcriptomes reveals a transcription regulatory network encompassing biosynthetic gene clusters. Adv. Sci. 11, 2403912 (2024). 10.1002/advs.202403912

[26] Jönsson, M. et al. Machine learning uncovers the transcriptional regulatory network for the production host *Streptomyces albidoflavus*. Cell Rep. 44, 115392 (2025). 10.1016/j.celrep.2025.115392

[27] Kwon, M. J. et al. Beyond the biosynthetic gene cluster paradigm: genome-wide coexpression networks connect clustered and unclustered transcription factors to secondary metabolic pathways. Microbiol. Spectr. 9, e00898–21 (2021). 10.1128/spectrum.00898-21

[28] Pinilla, L., Toro, L. F., Laing, E., Alzate, J. F. & Ríos-Estepa, R. Comparative transcriptome analysis of *Streptomyces clavuligerus* in response to favorable and restrictive nutritional conditions. Antibiotics 8, 96 (2019). 10.3390/antibiotics8030096

[29] Wu, J. et al. Comparative transcriptome analysis demonstrates the positive effect of the cyclic AMP receptor protein Crp on daptomycin biosynthesis in *Streptomyces roseosporus*. Front. Bioeng. Biotechnol. 9, 618029 (2021). 10.3389/fbioe.2021.618029

[30] Gong, J. et al. Transcriptome profiles of *Streptomyces clavuligerus* strains producing different titers of clavulanic acid. Sci. Data 10, 804 (2023). 10.1038/s41597-023-02727-6

[31] Jørgensen, T. S. et al. A treasure trove of 1034 actinomycete genomes. Nucleic Acids Res. 52, 7487–7503 (2024). 10.1093/nar/gkae523

[32] Langfelder, P. & Horvath, S. WGCNA: an R package for weighted correlation network analysis. BMC Bioinformatics 9, 559 (2008). 10.1186/1471-2105-9-559

[33] Greenfest-Allen E, Cartailler JP, Magnuson MA, Stoeckert CJ. iterativeWGCNA: iterative refinement to improve module detection from WGCNA co-expression networks. bioRxiv (2017). 10.1101/234062

[34] Blin, K. et al. antiSMASH 7.0: new and improved predictions for detection, regulation, chemical structures and visualisation. Nucleic Acids Res. 51, W46–W50 (2023). 10.1093/nar/gkad344

[35] Medema, M. H. et al. Minimum information about a biosynthetic gene cluster. Nat. Chem. Biol. 11, 625–631 (2015). 10.1038/nchembio.1890

[36] Draisma, A. et al. BiG-SCAPE 2.0 and BiG-SLiCE 2.0: scalable, accurate and interactive sequence clustering of metabolic gene clusters. Nat. Commun. 17, 2000 (2026). 10.1038/s41467-026-68733-5

[37] Gilchrist, C. L. M. & Chooi, Y.-H. clinker & clustermap.js: automatic generation of gene cluster comparison figures. Bioinformatics 37, 2473–2475 (2021). 10.1093/bioinformatics/btab007

[38] Emms, D. M. & Kelly, S. OrthoFinder: phylogenetic orthology inference for comparative genomics. Genome Biol. 20, 238 (2019). 10.1186/s13059-019-1832-y

[39] Martín, J. F., Casqueiro, J. & Liras, P. Secretion systems for secondary metabolites: how producer cells send out messages of intercellular communication. Curr. Opin. Microbiol. 8, 282–293 (2005). 10.1016/j.mib.2005.04.009

[40] Huo, L., Rachid, S., Stadler, M., Wenzel, S. C. & Müller, R. Synthetic biotechnology to study and engineer ribosomal bottromycin biosynthesis. Chem. Biol. 19, 1278–1287 (2012). 10.1016/j.chembiol.2012.08.013

[41] Rees, D. C., Johnson, E. & Lewinson, O. ABC transporters: the power to change. Nat. Rev. Mol. Cell Biol. 10, 218–227 (2009). 10.1038/nrm2646

[42] Yin, S. et al. Improvement of oxytetracycline production mediated via cooperation of resistance genes in *Streptomyces rimosus*. Sci. China Life Sci. 60, 992–999 (2017). 10.1007/s11427-017-9121-4

[43] Martín, J. F. & Liras, P. Molecular mechanisms of phosphate sensing, transport and signalling in *Streptomyces* and related Actinobacteria. Int. J. Mol. Sci. 22, 1129 (2021). 10.3390/ijms22031129

[44] Musrrat, S. et al. Prophage activation: an in silico platform for identifying prophage regulatory elements to inform phage engineering against drug-resistant bacteria. Life 15, 1417 (2025). 10.3390/life15091417 https://doi.org/10.3390/life15091417.

[45] Blatch, G. L. & Lässle, M. The tetratricopeptide repeat: a structural motif mediating protein-protein interactions. BioEssays 21, 932–939 (1999). 10.1002/(SICI)1521-1878(199911)21:11<932::AID-BIES5>3.0.CO;2-N.

[46] Bibb, M. The regulation of antibiotic production in *Streptomyces coelicolor* A3(2). Microbiology 142, 1335–1344 (1996). 10.1099/13500872-142-6-1335

[47] Amos, G. C. A. et al. Comparative transcriptomics as a guide to natural product discovery and biosynthetic gene cluster functionality. Proc. Natl Acad. Sci. USA 114, E11121–E11130 (2017). 10.1073/pnas.1714381115

[48] Grosjean, N. et al. A hemoprotein with a zinc-mirror heme site ties heme availability to carbon metabolism in cyanobacteria. Nat. Commun. 15, 3167 (2024). 10.1038/s41467-024-47486-z

[49] Hwang, S. et al. System-level analysis of transcriptional and translational regulatory elements in *Streptomyces griseus*. Front. Bioeng. Biotechnol. 10, 844200 (2022). 10.3389/fbioe.2022.844200

[50] Bode, H. B., Bethe, B., Höfs, R. & Zeeck, A. Big effects from small changes: possible ways to explore nature’s chemical diversity. ChemBioChem 3, 619–627 (2002). 10.1002/1439-7633(20020703)3:7<619::AID-CBIC619>3.0.CO;2-9

[51] Bilyk, O., Sekurova, O. N., Zotchev, S. B. & Luzhetskyy, A. Cloning and heterologous expression of the grecocycline biosynthetic gene cluster. PLoS ONE 11 (2016). 10.1371/journal.pone.0158682

[52] Donadio, S., Monciardini, P. & Sosio, M. Approaches to discovering novel antibacterial and antifungal agents. Methods Enzymol. 458, 3–28 (2009). 10.1016/S0076-6879(09)04801-0

[53] Kieser, T., Bibb, M. J., Buttner, M. J., Chater, K. F. & Hopwood, D. A. Practical Streptomyces Genetics. The John Innes Foundation (2000).

[54] Sánchez, S. et al. Carbon source regulation of antibiotic production. J. Antibiot. 63, 442–459 (2010). 10.1038/ja.2010.78

[55] Sánchez-Hidalgo, M. et al. Insights into the biosynthesis of kibdelomycin and amycolamicin from comparative biosynthetic gene cluster analysis and precursor incorporation studies. J. Nat. Prod. 88, 1988–1999 (2025). 10.1021/acs.jnatprod.5c00726

[56] Martin, M. Cutadapt removes adapter sequences from high-throughput sequencing reads. EMBnet.journal 17, 10–12 (2011). 10.14806/ej.17.1.200

[57] Langmead, B., Trapnell, C., Pop, M. & Salzberg, S. L. Ultrafast and memory-efficient alignment of short DNA sequences to the human genome. Genome Biol. 10, R25 (2009). 10.1186/gb-2009-10-3-r25

[58] Babraham Bioinformatics. FastQC: A quality control tool for high-throughput sequence data. Available from: https://www.bioinformatics.babraham.ac.uk/projects/fastqc/

[59] Wang, L., Wang, S. & Li, W. RSeQC: quality control of RNA-seq experiments. Bioinformatics 28, 2184–2185 (2012). 10.1093/bioinformatics/bts356

[60] Liao, Y., Smyth, G. K. & Shi, W. featureCounts: an efficient general purpose program for assigning sequence reads to genomic features. Bioinformatics 30, 923–930 (2014). 10.1093/BIOINFORMATICS/BTT656

[61] Ewels, P., Magnusson, M., Lundin, S. & Käller, M. MultiQC: summarize analysis results for multiple tools and samples in a single report. Bioinformatics 32, 3047–3048 (2016). 10.1093/bioinformatics/btw354

[62] Booth, T. J., Shaw, S., Cruz-Morales, P. & Weber, T. getphylo: rapid and automatic generation of multi-locus phylogenetic trees. BMC Bioinformatics 26, 21 (2025). 10.1186/S12859-025-06035-1.

[63] Cantalapiedra, C. P., Hernández-Plaza, A., Letunic, I., Bork, P. & Huerta-Cepas, J. eggNOG-mapper v2: functional annotation, orthology assignments, and domain prediction at the metagenomic scale. Mol. Biol. Evol. 38, 5825–5829 (2021)10.1093/molbev/msab293.

[64] Huerta-Cepas, J. et al. eggNOG 5.0: a hierarchical, functionally and phylogenetically annotated orthology resource based on 5090 organisms and 2502 viruses. Nucleic Acids Res. 47, D309–D314 (2019) 10.1093/nar/gky1085

[65] Tatusov, R. L., Galperin, M. Y., Natale, D. A. & Koonin, E. V. The COG database: a tool for genome-scale analysis of protein functions and evolution. Nucleic Acids Res. 28, 33–36 (2000). 10.1093/nar/28.1.33

[66] Rezaie, N., Reese, F. & Mortazavi, A. PyWGCNA: a Python package for weighted gene co-expression network analysis. Bioinformatics 39, btad415 (2023). 10.1093/bioinformatics/btad415

[67] Levassor, L. et al. StreptoCAD: an open-source software toolbox automating genome engineering workflows in streptomycetes. ACS Synth. Biol. 14, 4355–4372 (2025). 10.1021/ACSSYNBIO.5C00261

[68] Virtanen, P. et al. SciPy 1.0: fundamental algorithms for scientific computing in Python. Nat. Methods 17, 261–272 (2020). 10.1038/s41592-019-0686-2

[69] Benjamini, Y. & Hochberg, Y. Controlling the false discovery rate: a practical and powerful approach to multiple testing. J. R. Stat. Soc. B 57, 289–300 (1995). 10.1111/j.2517-6161.1995.tb02031.x

