## Supplementary_File for "Toward understanding drivers of specialized metabolism in Actinomycetota: insights from 1432 transcriptomics datasets for 132 strains"

*Corresponding author

Contact Information:

### Supplementary Results

#### Identification of the BGC that encodes the red compound in *Streptomyces sp.* NBC_00906

Region 14 (antiSMASH v7.0) was identified as the likely BGC encoding the red pigment in *Streptomyces sp.* NBC_00906 because it encoded a type II PKS cluster, which frequently produce highly conjugated compounds, and its expression pattern in the RNA-seq data correlated with the presence of the red molecule across the eight media. To confirm this link, we inactivated a core gene within the cluster encoding a β-ketoacyl-ACP synthase (KSα) using CRISPR base editing. Inactivation of this gene (locus ID: OG694_25700) resulted in complete loss of red pigmentation in cultures grown in DNPM medium compared to the WT strain (Supplementary Figure 5A). Consistent with this phenotype, HPLC analysis of culture extracts from the knockout strains at 1.30 min and 2.85 min, the retention times of red compound, showed the disappearance of chromatographic peaks with the characteristic UV pattern of the compound. Wild-type extracts displayed characteristic absorbance maxima (λmax) at approximately 435 nm and 547–548 nm, whereas these spectral features were strongly reduced or absent in the knockout strain (Supplementary Figure 5C,D).

### Supplementary Methods

#### Construction of BGC knockout mutant

CRISPR base editing (CRISPR-BEST) was used to inactivate the BGC responsible for red compound production by introducing premature STOP codons at defined positions in core biosynthetic genes [1]. Spacer sequences were designed using CRISPy-web [2], and CRISPR-BEST plasmids were constructed following a previously described protocol [1]. Spacers were integrated into the CRISPR-BEST plasmid backbone using an ssDNA oligo-bridging strategy.

The following procedure was used for construction of the CRISPR-BEST plasmids. Plasmid backbones were digested with NcoI at 37°C for 1 h and subsequently dephosphorylated using FastAP at 37°C for 30 min, followed by enzyme inactivation at 65°C for 10 min. Spacer oligonucleotides (20 nt) with 20 nt overlaps to the plasmid backbone were ordered from Integrated DNA Technologies (IDT, Coralville, USA). Two spacer sequences were evaluated in this study: Spacer01 (CCAGTGCCGCAGGCTCGACA) and Spacer02 (GCCAGTGC-CGCAGGCTCGAC), yielding plasmids pL000 and pL0002 respectively by use of AR01 (CCGGTTGGTAGGATCGACGGCCAGTGCCGCAGGCTCGACAGTTTTAGAGCTAGAAATAGC) and AR02 (CCGGTTGGTAGGATCGACGGGCCAGTGCCGCAGGCTCGACGTTTTAGAGCTAGAAATAGC) primers.

Correct assembly of the constructed plasmids was verified by Sanger sequencing using primers AR03 (TGTGTGGAATTGTGAGCGGATA) and AR04 (CCCATTCAAGAACAGCAAGCAG). Sequence-verified plasmids were introduced into Escherichia coli ET12567 (pUB307) by electroporation. Apramycin-resistant ET12567 transformants were subsequently used for intergeneric conjugation with

Streptomyces NBC 00906, as described above. Apramycin-resistant exconjugants were selected and screened by colony PCR.

Colony PCR was performed using primer pairs AR05 (CAGGTGCGGGGACAGGTACG) and AR06 (CAGGTGCGGGGACAGGTACG) for checking stop codon introduction and Sanger sequencing of the edited gene. PCR reactions were carried out using Q5 High-Fidelity Master Mix (New England Biolabs; M0492S) according to the manufacturer's instructions. For colony PCR, single colonies were collected directly from agar plates using sterile toothpicks and resuspended in 50 µL dimethyl sulfoxide (DMSO). Cell suspensions were subjected to two cycles of boiling and freezing (15 min each), and 1 µL of the treated suspension was used directly as template DNA. PCR products were analysed by Sanger sequencing using the corresponding screening primers to confirm the presence of the intended mutations.

Among the two constructs tested, only introduction of pL0001 resulted in successful generation of the desired premature STOP codon (Q66 STOP) in the targeted T2PKS biosynthetic gene (locus ID: OG694 25700). An additional CRISPR base-editing attempt targeting a second biosynthetic gene (locus ID: OG694 25695) did not result in successful introduction of a premature STOP codon.

**HPLC–DAD analysis of wild-type and knockout extracts**

To compare metabolite profiles between the WT and knockout (KO) strains, cultures were centrifuged to remove solid material and the resulting supernatants were transferred to fresh centrifuge tubes. Clarified supernatants were directly injected into an UltiMate 3000 HPLC system (Thermo Fisher Scientific) equipped with an Agilent Eclipse Plus C18 column (4.6 × 100 mm, 3 μm). Solvent A consisted of water supplemented with 0.1% formic acid, and solvent B was acetonitrile. The chromatographic method was as follows: isocratic elution at 20% B for 1 min; a linear gradient from 20% to 40% B between 1 and 2 min; isocratic elution at 40% B from 2 to 5.5 min; a linear gradient from 40% to 90% B between 5.5 and 7.5 min; and isocratic elution at 90% B from 7.5 to 9 min. Prior to each injection, the column was equilibrated for 3 min at 20% B. The flow rate was maintained at 1 mL min−1, and the column oven temperature was set to 30◦C. Detection was performed using a diode array detector operated in three-dimensional (3D) mode, enabling simultaneous acquisition of UV chromatograms and UV–Vis spectra.

### Supplementary Figures

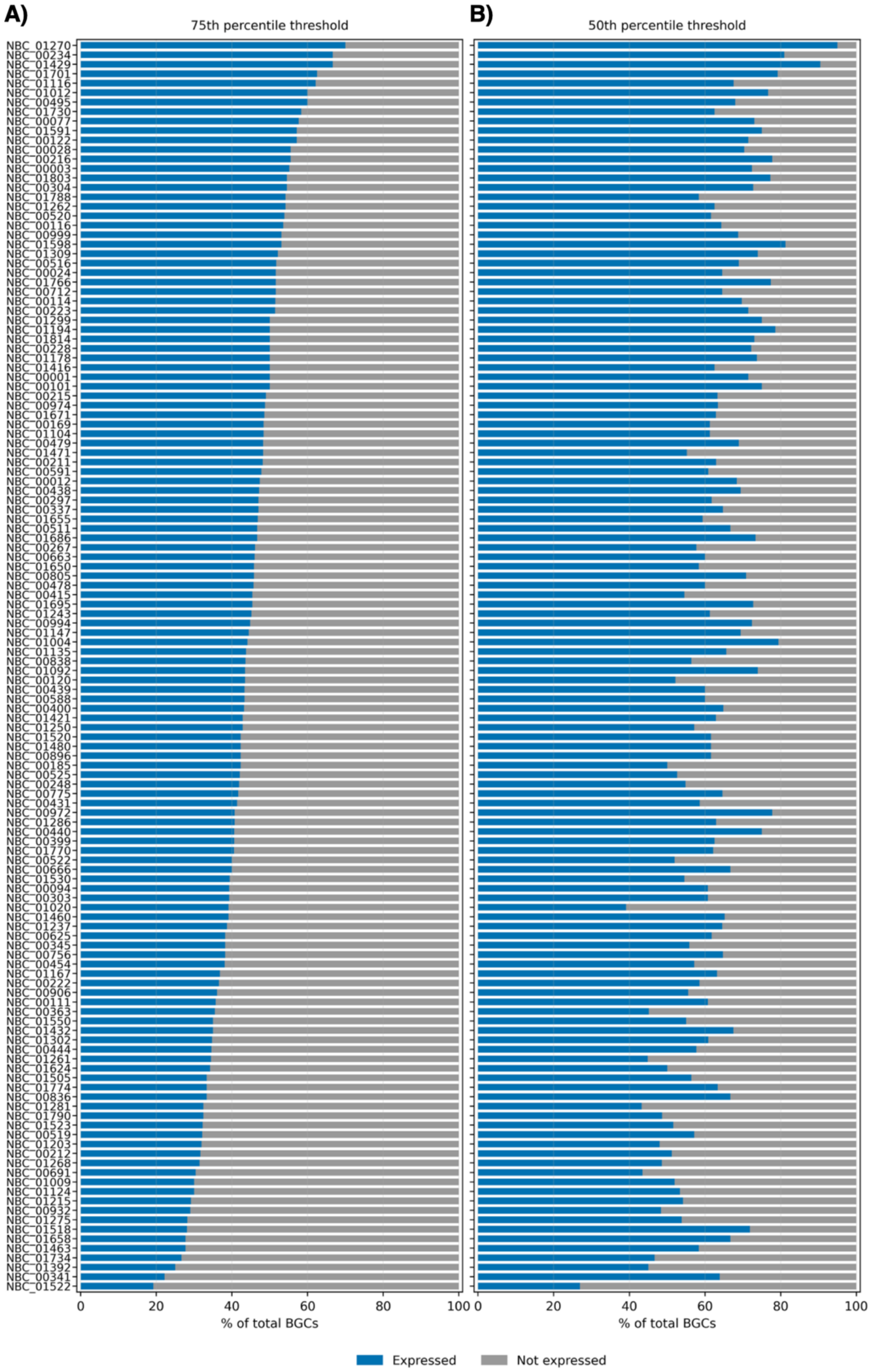

**Fig. 1** Fraction of expressed versus silent BGCs per strain. Horizontal bars show the proportion of BGCs in each genome categorized as expressed (blue) and not expressed (gray). A) 75th-percentile threshold. B) 50th-percentile threshold (same order as panel A).

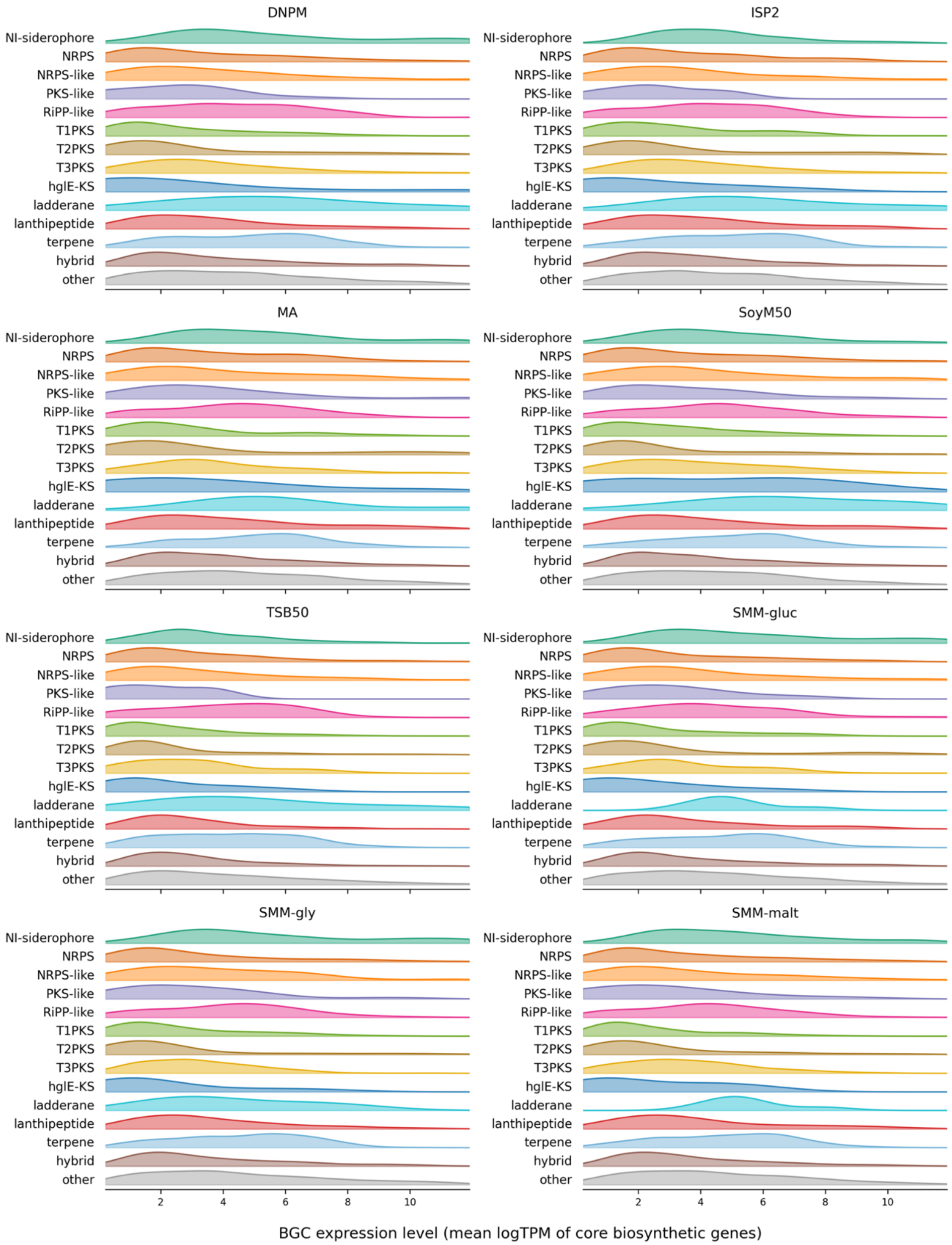

**Fig. 2**. Distribution of BGC expression levels across media and BGC types. Ridgeline density plots show the distribution of BGC expression levels for each BGC type across growth media. Each BGC was assigned one expression value per medium, calculated as the mean logTPM of its core biosynthetic genes. Colors indicate BGC type.

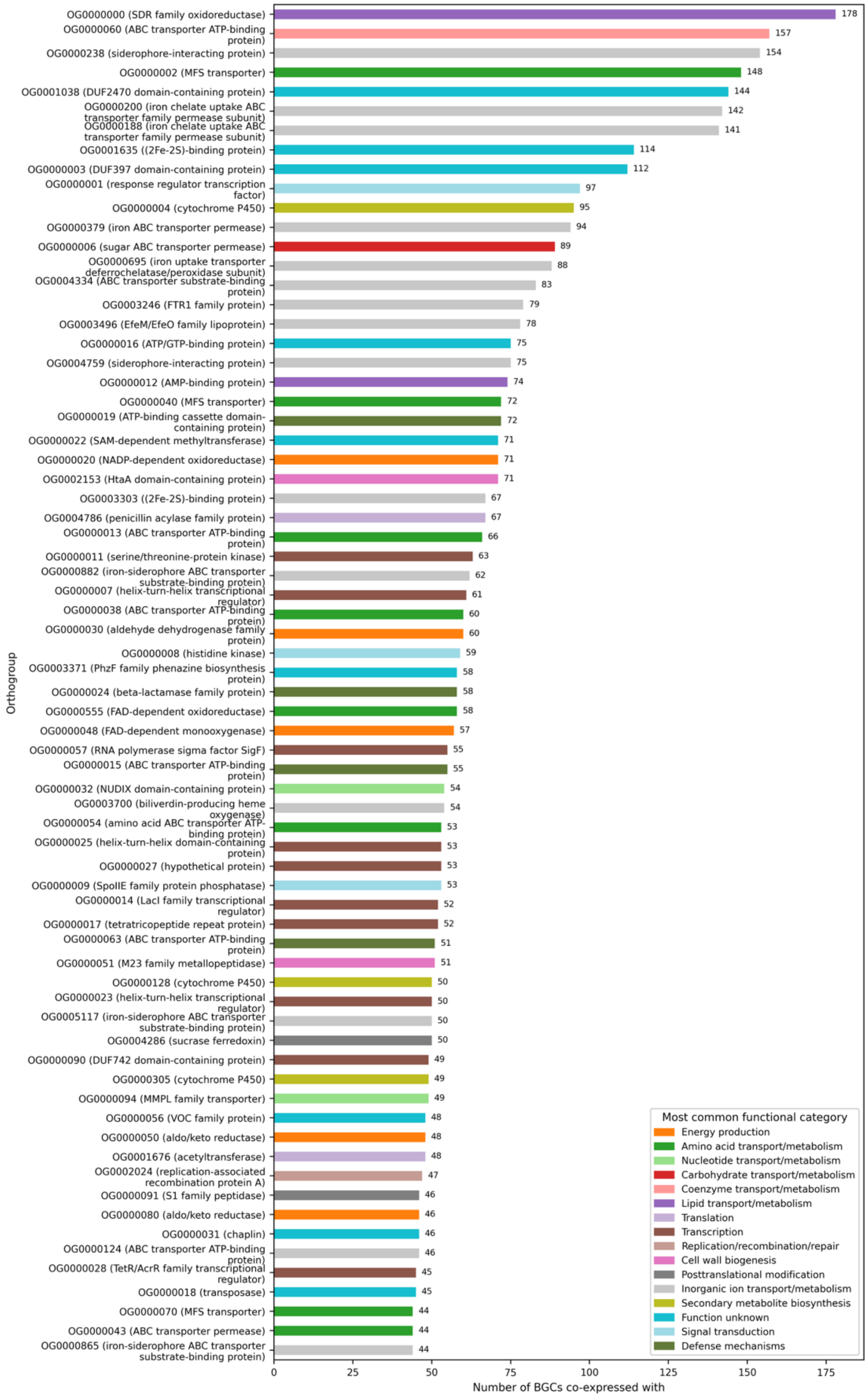

**Fig. 3**. Orthogroups frequently associated with BGC expression. Horizontal bar plot showing the top 70 orthogroups ranked by the number of distinct biosynthetic BGCs with which they are co-expressed. Each orthogroup is labeled with its identifier and the most frequent product annotation observed among its member genes. Bars are colored by the most common COG functional category within each orthogroup.

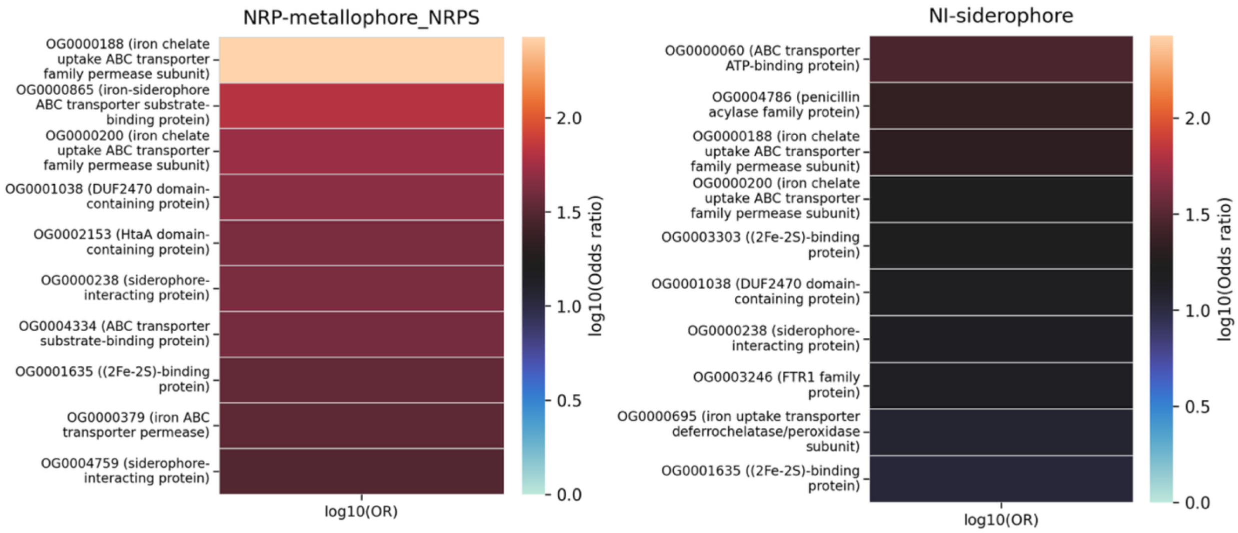

**Fig. 4** NRP-metallophore-NRPS and NI-siderophore BGC-type orthogrouop co-expression enrichment. Heatmaps show the top orthogroups most strongly enriched in co-expression with each BGC type, quantified as log10 odds ratios. Rows represent individual orthogroups, and colors indicate the strength of enrichment. In parentheses next to each orthogroups the most common gene product of the orthogroup is shown. Only the top 10 enriched genes per BGC type are shown.

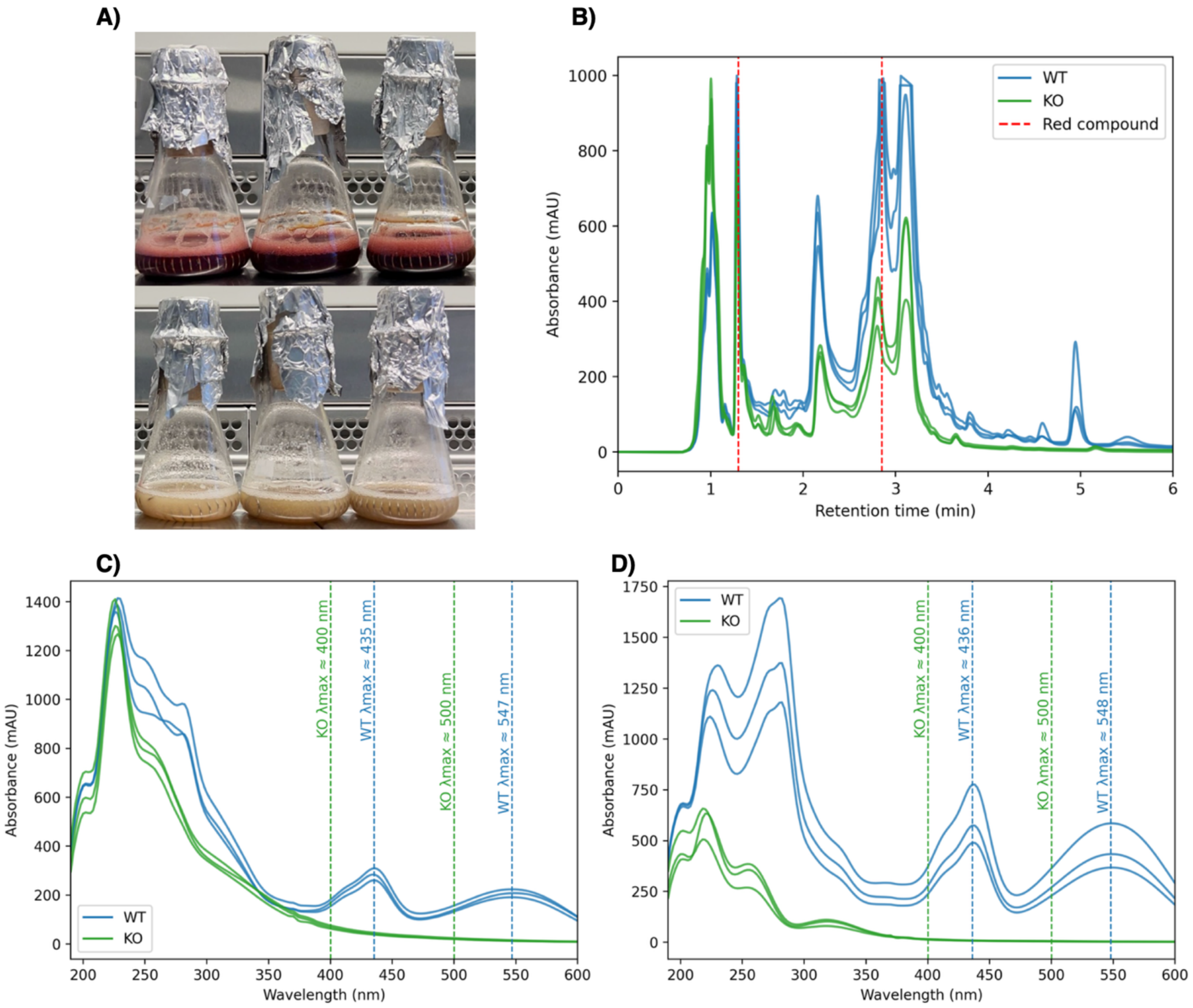

**Fig. 5** Genetic validation of the BGC responsible for red molecule production in *Streptomyces sp.* NBC_00906. A) Phenotypic comparison of the wild-type (WT) strain (top) and the CRISPR base-edited knockout (KO) strain (bottom) grown in DNPM medium, showing complete loss of red pigmentation upon disruption of a core T2PKS gene. B) UV chromatograms recorded at 257 nm (0–6 min) for WT and KO culture extracts. Dashed red lines indicate the retention times (~1.30 min and ~2.85 min) corresponding to the red compound. C) UV-Vis spectra extracted at RT ≈ 1.30 min from WT and KO chromatographic peaks. D) UV–Vis spectra extracted at RT ≈ 2.85 min from WT and KO chromatographic peaks. At both timepoints, the characteristic maxima of the red compound at 437 and 547 nm are absent in the KO samples. Spectra represent biological replicates (n = 3 per condition).

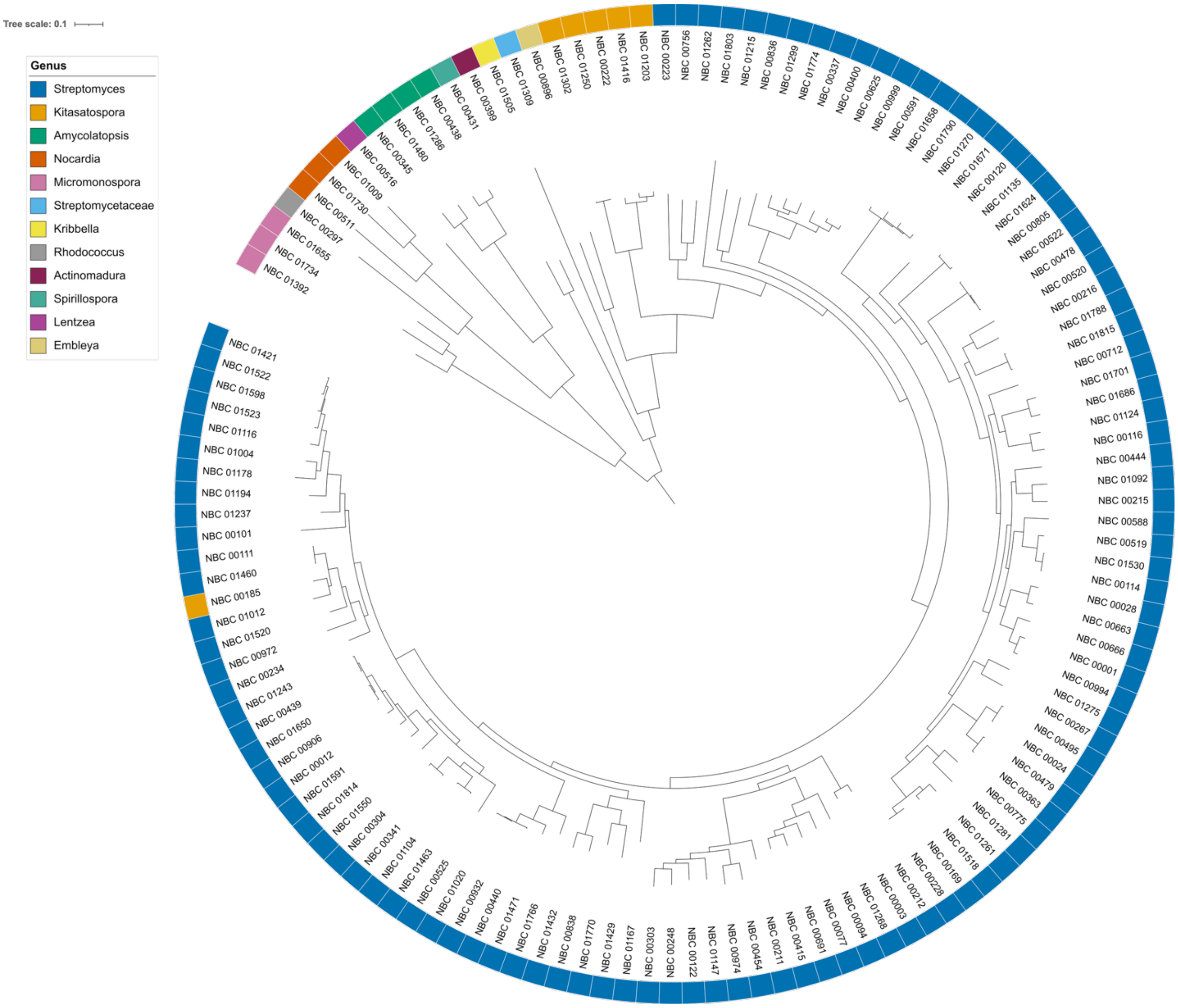

**Fig. 6** Phylogenetic distribution of the 132 Actinomycetota strains. The tree includes 132 strains and was midpoint-rooted for visualization. Tips are labelled by NBC strain identifier, and the outer color strip indicates genus. Branch lengths represent inferred phylogenetic distances.

**Supplementary References**

[1] Tong, Y. et al. Highly efficient DSB-free base editing for streptomycetes with CRISPR-BEST. *Proc. Natl Acad. Sci. USA* 116, 20366–20375 (2019). https://doi.org/10.1073/pnas.1913493116

[2] Blin, K., Shaw, S., Tong, Y. & Weber, T. Designing sgRNAs for CRISPR-BEST base editing applications with CRISPy-web 2.0. *Synth. Syst. Biotechnol.* 5, 99–102 (2020). https://doi.org/10.1016/j.synbio.2020.05.005
